# Mosaic origin of the eukaryotic kinetochore

**DOI:** 10.1101/514885

**Authors:** Jolien J.E. van Hooff, Eelco Tromer, Geert J.P.L. Kops, Berend Snel

## Abstract

The emergence of eukaryotes from ancient prokaryotic lineages was accompanied by a remarkable increase in cellular complexity. While prokaryotes use simple systems to connect DNA to the segregation machinery during cell division, eukaryotes use a highly complex protein assembly known as the kinetochore. Although conceptually similar, prokaryotic segregation systems and eukaryotic kinetochore proteins share no homology, raising the question of the origins of the latter. Using large-scale gene family reconstruction, sensitive profile-versus-profile homology detection and protein structural comparisons, we here reveal that the kinetochore of the last eukaryotic common ancestor (LECA) consisted of 52 proteins that share deep evolutionary histories with proteins involved in a few prokaryotic processes and a multitude of eukaryotic processes, including ubiquitination, chromatin regulation and flagellar as well as vesicular transport systems. We find that gene duplications played a major role in shaping the kinetochore: roughly half of LECA kinetochore proteins have other kinetochore proteins as closest homologs. Some of these (e.g. subunits of the Mis12 complex) have no detectable homology to any other eukaryotic protein, suggesting they arose as kinetochore-specific proteins de novo before LECA. We propose that the primordial kinetochore evolved from proteins involved in various (pre-)eukaryotic systems as well as novel proteins, after which a subset duplicated to give rise to the complex kinetochore of LECA.

## Introduction

During cell division, eukaryotes divide their duplicated chromosomes over both daughter cells by means of a microtubule-based apparatus called the spindle. Central to this process are kinetochores; large multi-protein structures that are built upon centromeric DNA and that connect chromosomes to microtubules. Although species vary hugely in how they exactly coordinate and execute chromosome segregation [1–4], all eukaryotes use a microtubule-based spindle apparatus, and therefore the last eukaryotic common ancestor (LECA, Figure 1A) likely laboured one as well. Consequently, LECA’s chromosomes probably contained a centromere and assembled a kinetochore. While the centromeric DNA sequences of current-day eukaryotes are strikingly different between species and too diverse to reconstruct LECA’s centromeric DNA [5], their proteomes did allow for the inference of LECA’s kinetochore. In previous work, we found that the LECA kinetochore was a complex structure, consisting of at least 49 different proteins [6].

**Figure 1.**
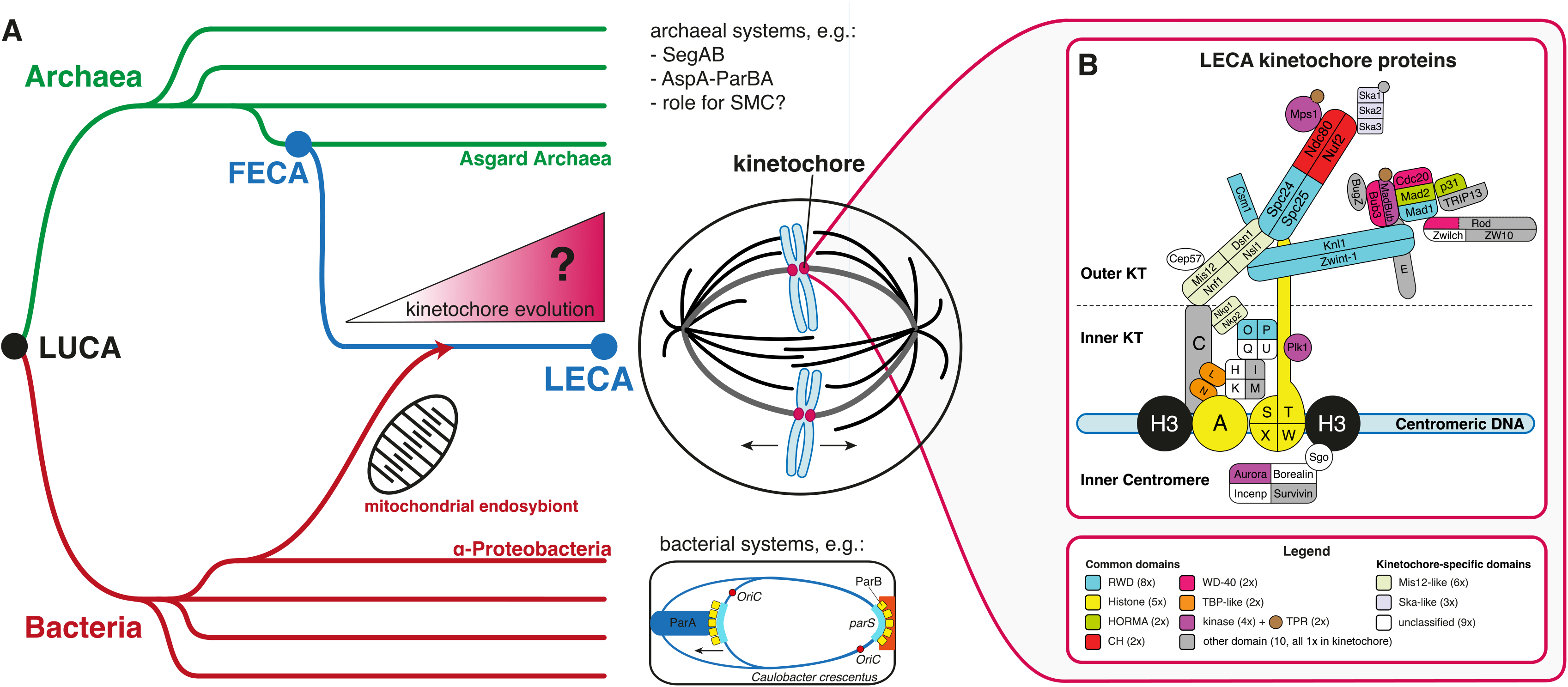
The eukaryotic kinetochore and mitotic machinery originated between FECA and LECA. (**A**) How did the eukaryotic kinetochore originate and evolve between FECA and LECA? Eukaryotes (blue) descended from Archaea (green), and are likely closely related to the Asgard superphylum [53]. This Asgard-related lineage incorporated an alphaproteobacterium via endosymbiosis; the latter gave rise to the eukaryotic mitochondrion. As far as currently characterized, Archaea and Bacteria (red) do not separate their duplicated chromosome(s) via a mitotic spindle [7–9]. For example, bacteria such as *Caulobacter crescentus* operate the *parABS* partitioning system, in which *parS* sites are recognized by the protein ParB, stimulating ParA, which in turn pulls or pushes the chromosomes apart [8]. Due to these differences, the mitotic spindle and the kinetochore probably originated between the first eukaryotic common ancestor (FECA) and the last eukaryotic common ancestor (LECA). LUCA: last universal common ancestor. (**B**) The kinetochore of LECA consisted of 52 proteins that contain domains found in other, non-kinetochore eukaryotic proteins as well (‘common domains’), or that are unique to the kinetochore (‘kinetochore-specific domains’). Proteins were inferred to have been part of the LECA kinetochore as described in SI. KT: kinetochore.

The LECA kinetochore was not directly derived from a prokaryotes, because prokaryotes employ protein assemblies that are not homologous to the eukaryotic kinetochore to link their DNA to the segregation machinery [7–9] (Figure 1A). Like many other unique eukaryotic cellular systems, the LECA kinetochore must thus have originated after the first eukaryotic common ancestor (FECA) diverged from prokaryotes. Between FECA and LECA, the pre-eukaryotic lineage evolved from relatively simple and small prokaryotic cells to complex, organelle-bearing cells that are organized in a fundamentally different manner, a process referred to as ‘eukaryogenesis’. What evolutionary events underlie eukaryogenesis is a major question [10], to which answers are offered by investigations into specific eukaryotic systems [11]. Studies on for example the spliceosome, the intracellular membrane system and the nuclear pore revealed that (repurposed) prokaryotic genes played a role in their origin, as did novel, eukaryote-specific genes and gene duplications, albeit in varying degrees and in different manners [12–14].

In this study, we address the question how the kinetochore originated. Leveraging the power of detailed phylogenetic analyses, improved sensitive sequence searches and novel structural insights, we traced the evolutionary origins of the 52 proteins we now assign to the LECA kinetochore. Based on our findings, we propose that the LECA kinetochore is of mosaic origin: it contains proteins that share ancestry with proteins involved in various core eukaryotic processes as well as completely novel proteins. After recruitment, many of these proteins duplicated, accounting for a 50% increase in kinetochore extent and thereby for the complex LECA kinetochore.

## Results

### The LECA kinetochore

To study how the LECA kinetochore originated, we first determined its protein content. For each protein present in current-day human and yeast kinetochores, we asked whether A) it was encoded in the genome of LECA, based on its distribution in current-day eukaryotes and B) whether it likely operated in the LECA kinetochore, based on functional information from current-day species. We inferred a protein to have been encoded by the LECA genome if it is found in both Opimoda and Diphoda, whose divergence likely represents the root of the eukaryotic tree of life (Figure S4, SI Text). We here extend our previous analyses [6] with orthologous groups of Nkp1, Nkp2 and Csm1 (see for further discussion SI Text, Figure 4A). Altogether, we propose that the LECA kinetochore consisted of at least 52 proteins (Figure 1B, Table S1). Of note: our reconstruction confirms [6] that most of the CCAN proteins (Constitutive Centromere Associated Network proteins) were part of the LECA kinetochore (SI Text).

### Identifying ancient homologs of kinetochore proteins

In order to elucidate the ancient, pre-LECA homologs (either eukaryotic or prokaryotic) of LECA kinetochore proteins, we applied sensitive profile-versus-profile homology searches (Table S2), followed by phylogenetic tree constructions (Figures S1, S3A), or, if available, published phylogenetic tree interpretations (SI Text). If literature and/or structural studies provided additional information on ancient relationships, we also included these as evidence for a homologous relationship of a kinetochore protein (Table S3). For each LECA kinetochore protein, we examined which proteins comprise its closest homologs before LECA (Table 1). These proteins were classified as eukaryotic or prokaryotic, and as kinetochore or non-kinetochore (SI Data and Methods). In order to allow different domains in a single protein to have different evolutionary histories, we primarily searched for homologs on the domain level, and represent these as a single ‘domain’ in Table 1 if they share their evolutionary history as part of a single protein.

**Table 1.**
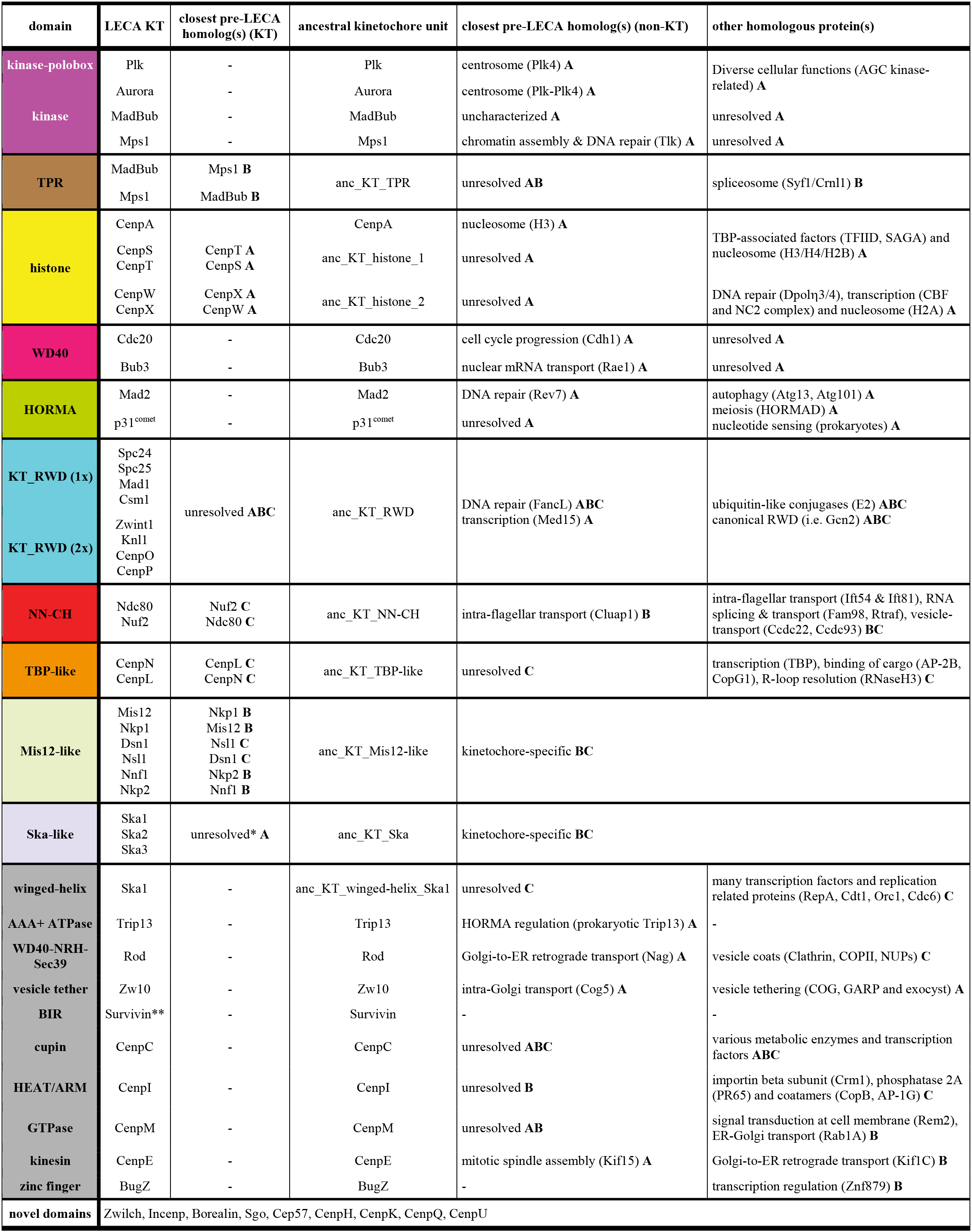
Ancient homologs of kinetochore (KT) domains and their functions (see Figure 1B, 5). Note that if multiple domains have a shared evolutionary history, we regard them as a single unit in this table (kinase-polo box, WD40-NRH-Sec39). Some domains were recruited to the kinetochore before they duplicated to give rise to multiple kinetochore proteins. Those initial kinetochore entities are the ‘ancestral kinetochore units’. If a protein does not have closely related homologs in the kinetochore, the protein itself was the ancestral unit that got involved in the kinetochore. For all relationships, we indicate which type of evidence we have for it. **A**: phylogenetic tree, **B**: hit in profile-profile search, **C**: structure and/or literature. *The phylogeny of Ska1, Ska2 and Ska3 cannot be rooted, therefore it is unknown which are the each other’s closest paralog. **The BIR domain is involved in multiple processes in animals, but the kinetochore (inner centromere) function might be the ancestral one, because this is also reported in budding and fission yeast, which only have one BIR domain protein.

We inferred the closest homologs of kinetochore proteins on the domain level (Table 1), using gene phylogenies for 19/55 domains (35%), profile-versus-profile searches for 5/55 (9%) and structural information for 6/55 (11%,). For ten others (18%), we used a combination. For a total of 40 domains we could identify the closest homolog. For six (11%) of the remaining ones, we found homologs but could not resolve which is closest, and for the other nine (16%) we could not find any ancient homologs at all (Table 1).

### Evolutionary histories of kinetochore proteins

Below we discuss the evolutionary history of LECA kinetochore proteins per protein domain, including their affiliations to other eukaryotic cellular processes, their prokaryotic homologs and their ancient duplications within the kinetochore (see Table 1 for overview).

#### Kinetochore RWDs

The RWD (RING-WD40-DEAD)-like domains in kinetochore proteins are highly diverged and non-catalytic members of the structural superfamily of E2 ubiquitin-like conjugases (UBC) of which both bacterial and archaeal homologs are involved in ubiquitin-like modification [15–17] (Figure 2, Table S3). For seven LECA kinetochore proteins the structure of their RWD domains were determined (Figure 2B). These form hetero- or homodimers with either a single RWD (Spc24-Spc25, Mad1-Mad1 and Csm1-Csm1) or double RWD configuration (CenpO-CenpP and Knl1). In contrast to previous efforts [15, 18], our sensitive profile-versus-profile searches now uncovered significant sequence similarity of the Knl1-binding protein Zwint-1 with other double RWDs, suggesting that Zwint-1 and Knl1 form an RWD heterodimer similar to CenpO-CenpP (SI Text, Figure S2). To examine the origins of both single and double kinetochore RWDs, we aligned archaeal, bacterial and eukaryotic UBC proteins and performed a phylogenetic analysis (SI Text & Data and Methods, Figures S1E, S3). We found that kinetochore RWDs and other RWDs are more closely related to each other (bootstrap:96/100) than to eukaryotic and archaeal E2-like conjugases (bootstrap:77/100). A single archaeal (Asgard) sequence clustered at the base of the canonical eukaryotic RWDs, suggesting that FECA may have already contained an RWD-like domain. As supported by our profile-versus-profile searches (Table S2) and structural alignments (Figure 2B, Table S3, File S151), most kinetochore RWDs are each other’s closest homologs, indicating that kinetochore RWDs in LECA arose from a single ancestor, which does not contain the canonical RWD domains. Possibly, this group also includes a single (Med15) and a double RWD protein (FancL). We were however not able to reliably reconstruct the exact order by which the kinetochore RWD proteins arose. We suggest that kinetochore RWDs and other RWDs (i.e. Gcn2, FancL and Rwd1-4), evolved from a non-catalytic E2 ubiquitin-like conjugase as part of an extensive radiation and neofunctionalization of the UBC family during eukaryogenesis (Figure 2B).

**Figure 2.**
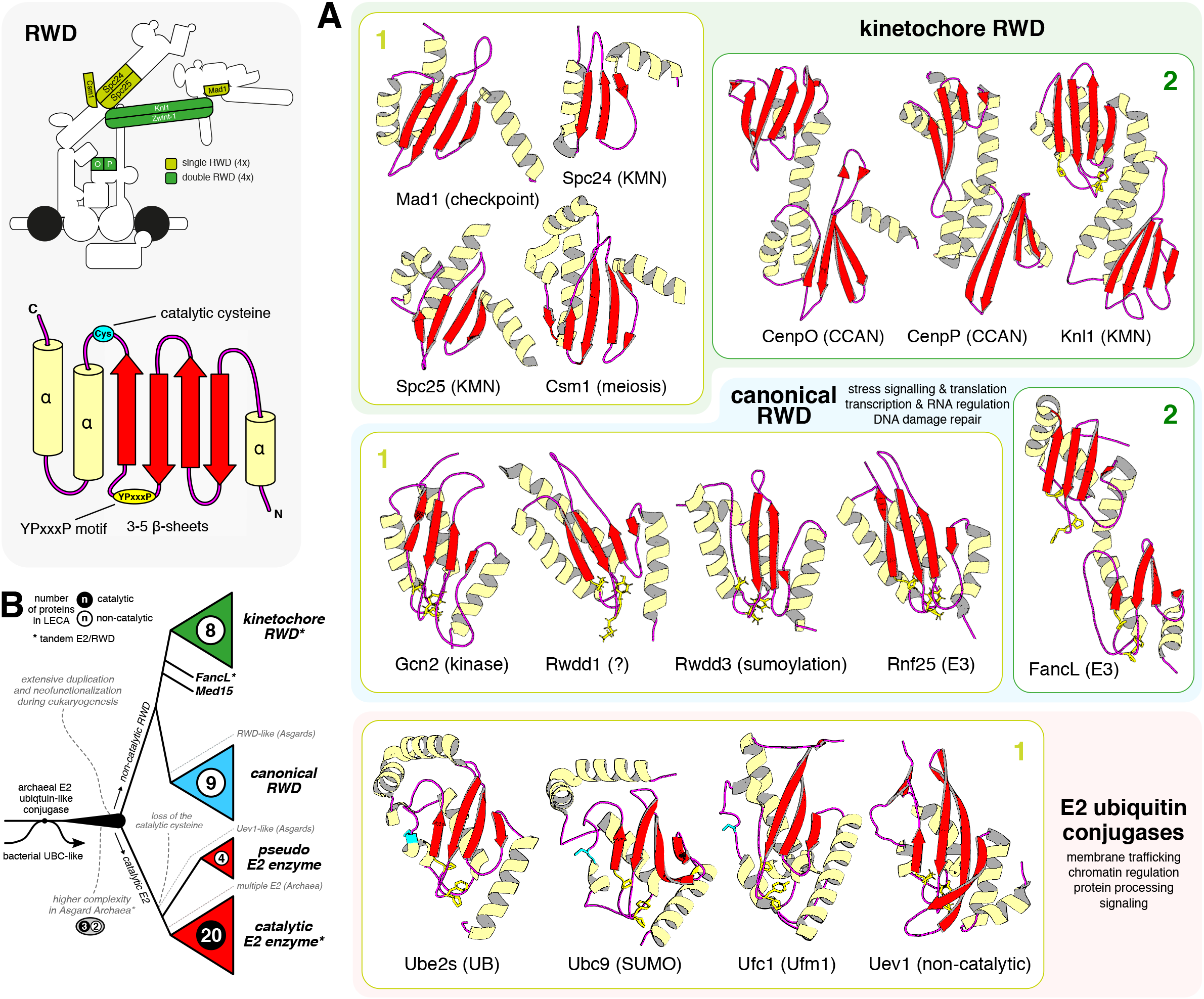
Kinetochore RWDs are an expanded class of non-catalytic E2 ubiquitin-like conjugases. Legend: (top) overview of the position of the eight kinetochore proteins with RWD domains. Kinetochore RWD proteins have a similar structural topology: N-terminal coiled-coil and a C-terminal single (light green) or a double (green) RWD. (bottom) secondary structure of E2 and RWD proteins of the UBC superfamily that is characterized by a ‘β-meander’ of 3-5 β-sheets, enclosed by α-helices at both termini, a ‘YPxxxP’ motif that often resides in between the third and the fourth β-sheet, and a cysteine residue involved in ubiquitination (lost in RWD). (**A**) The UBC superfamily consists of three distinct classes: (1) E2 ubiquitin conjugases, which function in ubiquitin-like modification and non-catalytic paralogs that interact with ubiquitin (Uev1), (2) canonical RWD proteins that operate as a dimerization domain to facilitate various E2/E3 ubiquitin-like ligation reactions (FancL-Ube2T and Rwdd3/Ubc9) and (3) RWD-like kinetochore proteins that form dimers and constitute the kinetochore superstructure (Spc24-Spc25, Knl1-Zwint-1 and CenpO-CenpP) and play a role in microtubule attachment regulation (Mad1 and Csm1). Per class, the structure of various members is depicted to show the overall structural and topological similarity and a known molecular function is indicated between brackets. If present, the YPxxxP (yellow) and the catalytic cysteine residues (cyan) are represented in the ‘sticks’ configuration. (**B**) A cartoon of the evolutionary reconstruction of the UBC superfamily (for annotated phylogenetic trees, see Figure S1E, S3). In short, extensive duplication and neofunctionalization of an archaeal E2 ubiquitin-like conjugase gave rise to a large complexity of catalytic and non-catalytic E2/RWD proteins in LECA (see numbers per class). Possibly, part of the eukaryotic complexity was already present in FECA, since Asgard archaea contain multiple E2 conjugases in addition to a non-catalytic E2 (Uev1-like) and an RWD-like domain (Figure S3). Bacterial UBCs likely represent an ancient protein modification system that has been optimized in archaeal lineage that are closely associated with FECA, but lateral transfers from archaeal and eukaryotic lineages were also detected (Figure S3, SI text).

#### Histones

The LECA kinetochore contained five histone proteins: CenpA and the CenpS-X-T-W tetramer. From FECA to LECA, an archaeal-derived histone-like protein [19, 20] duplicated many times, giving rise to variants involved in all aspects of eukaryotic chromatin complexity (Figure 3A). CenpA is a centromere-specific histone H3 variant and resulted from an ancient duplication before LECA [6, 20]. We found that CenpS-X-T-W arose by two duplications: CenpS-T (bootstrap:99/100) and CenpX-W (bootstrap:77/100), indicating a likely co-duplication of the two subunits of an ancestral heterodimer (see SI Text, Methods, Figure S1I), Furthermore, CenpS-T was phylogenetically affiliated to H2B-H3-H4-TFĲD-SAGA-related histones, while CenpX-W clustered with H2A-CBF-NC2-DPOE-Taf11-related histones (Figure 3A, Figure S1I). These affiliations in combination with an additional role of the CenpS-X dimer in the Fanconi anemia pathway [21, 22] signify that the origin of CenpS-X-T-W is interlinked with the emergence of the intricate eukaryotic transcription and DNA repair machinery.

**Figure 3.**
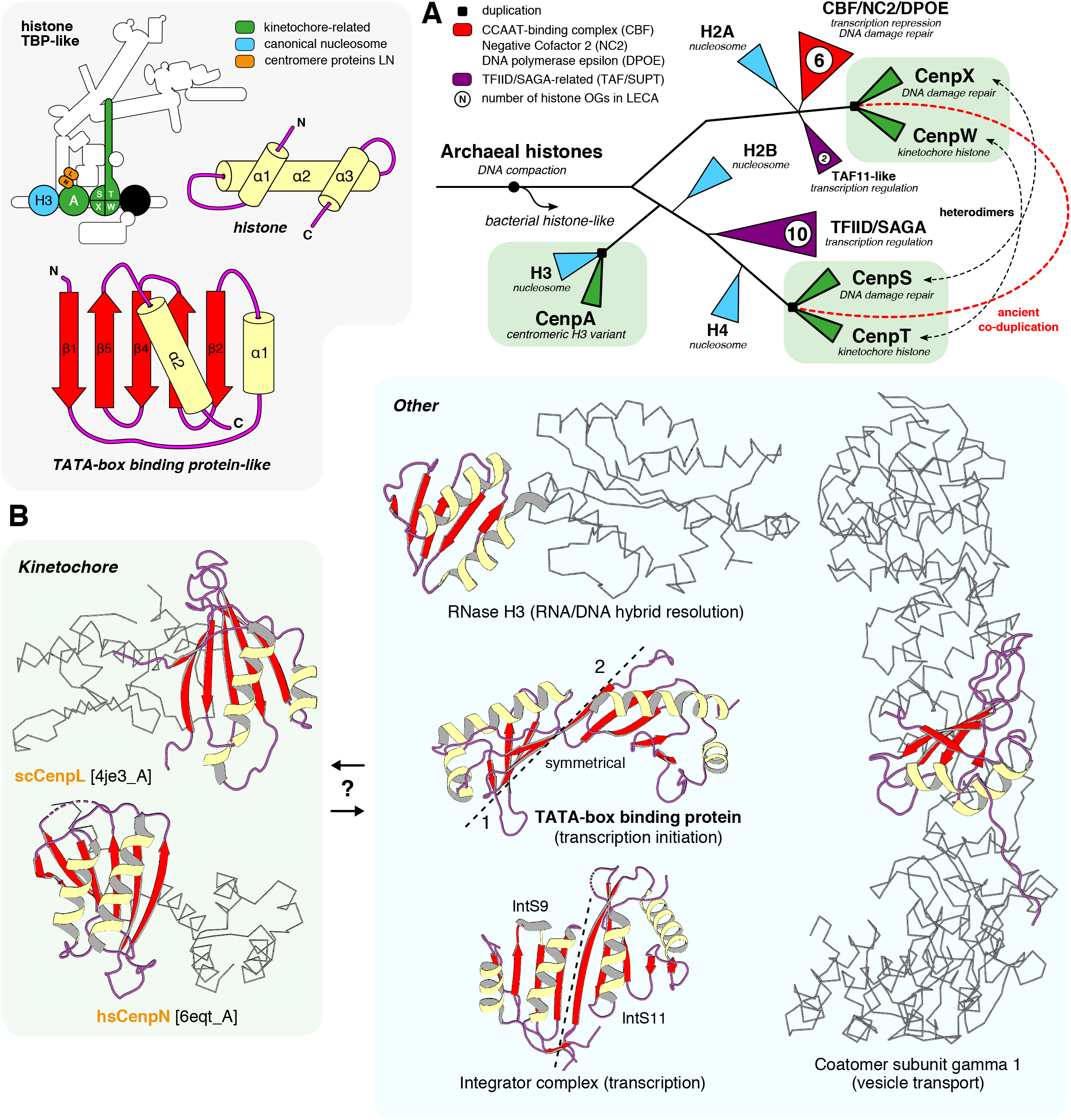
A common origin kinetochore histones and TBP-like proteins with complexes involved in DNA damage repair and transcriptional regulation. Legend: (top) an overview of the position of CenpA and CenpS-X-T-W (histones, green) and CenpL-N (TBP-like, orange) in the kinetochore. (bottom) histones consists of 3 α-helices with conserved interaction loops; the TBP-like fold is an elaborate set of curved β-strands that form an interaction surface for substrates, such as DNA, RNA and various protein motifs, but also constitute a possible dimer interface, resulting in an even larger extended β-sheet configuration. (**A**) A cartoon of the evolutionary reconstruction of kinetochore-related histone proteins CenpA and CenpS-T-X-W (Figure S1I). A histone of archaeal descent duplicated and subfunctionalized many times, giving rise to a large diversity of histone proteins in eukaryotes, including those involved in chromatin structure (nucleosome), transcriptional regulation (TAF/SUPT/NC2/CBF) and DNA damage repair (DPOE). Bacterial histone-like proteins were likely laterally transferred from either Archaea and/or eukaryotes (SI Text). CenpA is the closest homolog (paralog) of the nucleosomal histone H3. CenpS-T and CenpX-W, are likely each other’s closest paralogs, signifying a co-duplication of an ancient dimer to form the tetramer CenpS-X-T-W. The CenpS-X dimer also plays a role in the Fanconi Anemia pathway (DNA repair). OG: orthologous group. (**B**) CenpL and -N contain a TATA-box binding protein (TBP)-like fold. The degree of structural similarity between TBP-like proteins does not clearly indicate how CenpL-N are evolutionary related to other TBP-like domains. Yellow (helices) and red (sheets) show the location of a TBP-like domain in a subset of available TBP-like protein structures, the grey-ribbon representation indicates the non-homologous parts of the proteins; their cellular function is indicated between brackets. For CenpL-N, the used PDB accessions are shown.

#### TBP-like

CenpN and CenpL harbour a fold similar to the pseudo-symmetric DNA-binding domain of the TATA-box binding protein (TBP) [23–25]. Although we did not observe any significant sequence similarity for CenpL and CenpN (Table S2), we found structural similarity with a diverse group of proteins that function in nucleotide metabolism, in transcription and in vesicle transport [26] (Figure 3B, Table S3, File S152). TBP as well as various TBP-like DNA/RNA-related enzymes [26] were found in Archaea [27], suggesting eukaryotes acquired these proteins via vertical descent (Figure 1A). The structural similarities between CenpL-N and other TBP-like proteins did not indicate which are the most closely related. Nevertheless, given that they form a heterodimer [25], we propose that CenpL and CenpN are closest homologs, and that other TBP-like proteins are more distantly related.

#### Misl2-like

Through profile-versus-profile searches we discovered a previously hidden homology within the kinetochore: subunits of the Nkp complex were found to be homologous to subunits of the Mis12 complex. Combined with the similar structural topology of the Mis12 complex subunits, we infer that all subunits of these two complexes are homologous (Figure 4A, SI Text). We name these proteins Mis12-like. Sequence similarities indicated that Nnf1 and Nkp2 are most closely related to each other, as well as Mis12 and Nkp1, hence these pairs might result from the most recent duplications. Possibly, the Mis12 complex originated first, via intra-complex duplications, and subsequently the Nkp complex originated from co-duplication of the ancestors of Nnf1 and Mis12. We did not detect homologs of these Mis12-like proteins outside of the kinetochore.

**Figure 4.**
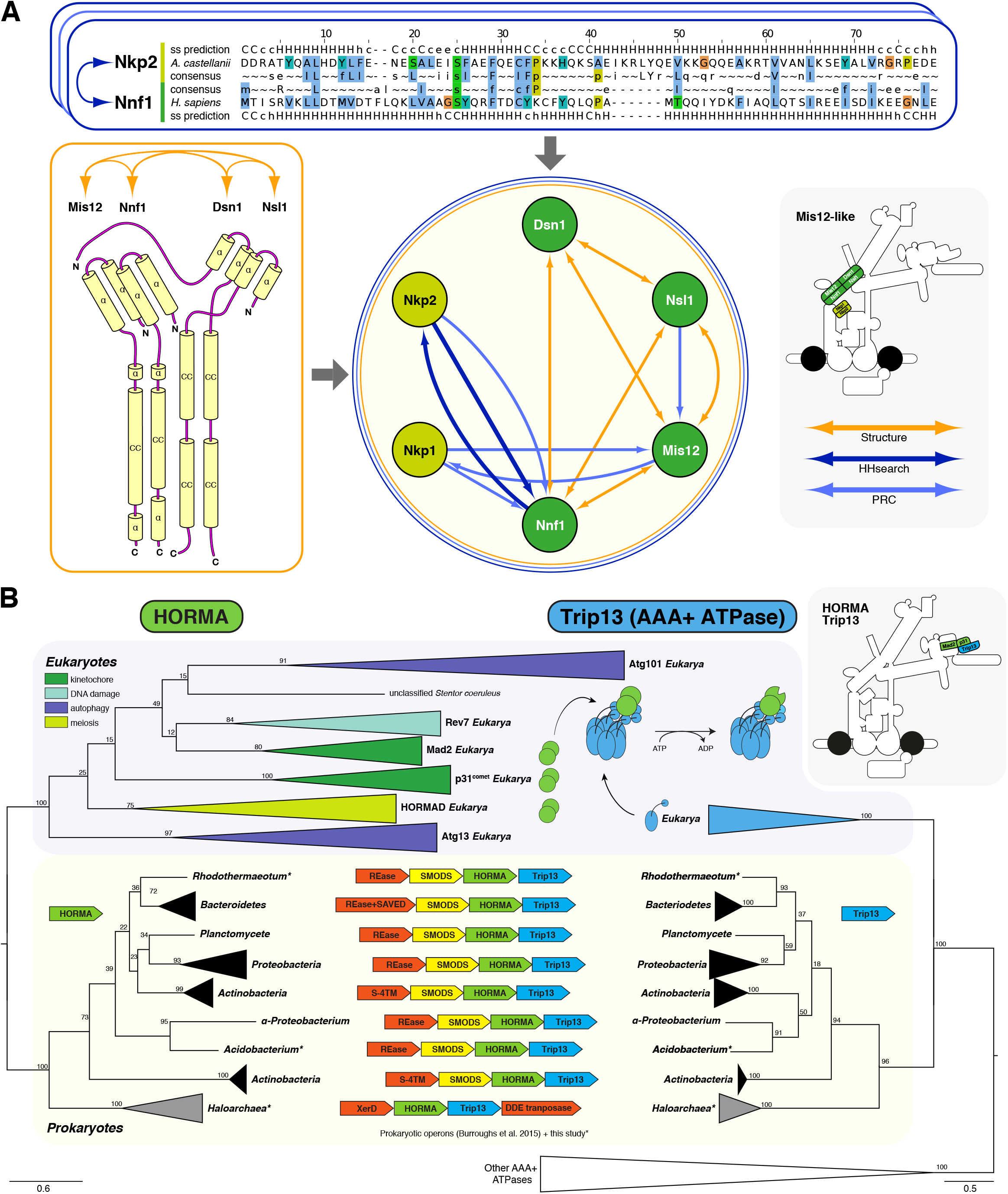
The Mis12 and Nkp complexes have a common ancestry and the HORMA-Trip13 module probably has a prokaryotic origin. (A) Profile-versus-profile hits (with HHsearch and PRC) and structural information [56, 57] indicate that all subunits of the Nkp and Mis12 complexes are homologous to each other (SI Text). Note that only one example of the profile-versus-profile hits is shown. Since Mis12 complex subunits are present across eukaryotes [6], we infer that also subunits of Nkp1 and Nkp2 were in LECA, as they resulted from pre-LECA duplications. Nkp2 and Nnf1 are each other’s best hit in profile-versus-profile searches, so possibly these proteins resulted from a relatively late duplication. The same holds for Nkp1-Mis12. (**B**) Phylogenetic trees of HORMA domain proteins and AAA+ ATPases. In eukaryotes, HORMAD and p31^comet^ are structurally modified by a Trip13 hexamer (upper panel, right side). These phylogenetic trees suggest that the eukaryotic HORMA domain and Trip13 were derived from prokaryotes. In prokaryotes, HORMA and Trip13 are present in a single operon, strongly suggesting that they also interact in these species and thus that this interaction is ancient. Moreover, this operon includes proteins that are involved in nucleotide signalling, suggesting prokaryotic HORMA and Trip13 are affiliated to this process [28]. The uncollapsed trees can be found in Figure S1F, S1G. Asterisks indicate the species for which we discovered a HORMA-Trip13 operon (see File S154 for annotation).

#### HORMA-Tripl3

Eukaryotic HORMA domain proteins operate in the kinetochore (Mad2, p31^comet^), autophagy (Atg13, Atg101), DNA repair (Rev7) and meiosis (HORMAD). The HORMA proteins p31^comet^ and HORMAD are structurally modified by Trip13, an AAA+ ATPase. Bacterial genomes also encode HORMA proteins and, interestingly, these co-occur in one operon with a Trip13-like AAA+ ATPase [28]. We additionally found the HORMA-Trip13-like operon in a few archaeal species that belong to the Haloarchaea (Figure 4B, File S154). The eukaryotic HORMA proteins are monophyletic, indicating FECA-to-LECA duplications (Figure S1F). Eukaryotic Trip13 sequences are more closely related to the prokaryotic Trip13-like sequences than to any other AAA+ ATPase (Figure S1G). How did eukaryotes acquire the HORMA-Trip13 module? While our phylogenetic analysis does not unequivocally indicate its ancestry, we propose that the pre-eukaryotic lineage derived the operon by horizontal transfer from Bacteria. Because in bacteria HORMA-Trip13 is part of operons with genes involved in nucleotide signalling [28], it might initially have fulfilled such a role in the pre-eukaryotic lineage, in which HORMA subsequently duplicated and neofunctionalized. As a result, HORMA-Trip13 got repurposed for eukaryote-specific processes, such as meiosis, autophagy and the kinetochore.

#### NN-Calponin Homology

CH (Calponin Homology) domain proteins operate in many different processes, including binding of actin and F-actin, and in various cellular signalling pathways [29]. In the kinetochore, they are the predominant microtubule-binding proteins. The ancestral function of this domain, which to our knowledge has not been found in prokaryotes, is not known. The kinetochore CH proteins seem to be part of a highly divergent subfamily of CH proteins (NN-CH) [30], which includes proteins involved in intraflagellar transport, ciliogenesis, the centrosome, vesicle-trafficking and possibly RNA transport [31–34]. It has been suggested this NN-CH subfamily is specialized towards binding microtubules, implying that the kinetochore function reflects the ancestral function [30].

#### Common eukaryotic domains: kinase, TPR, vesicle coats and tethers and WD40

In a detailed eukaryotic kinome phylogeny, the kinetochore kinases Plk and Aurora were closely related (Table 1, Figure S1D). The closest relative of Plk is Plk4, probably signalling an ancestral function for Plk in centrosome/basal body function, since Plk is also still found at the centrosome. Aurora diverged from a duplication prior to the Plk-Plk4 divergence, suggesting Plk and Aurora independently gained kinetochore functions after duplication. Alternatively, the Plk-Aurora ancestor operated in both the centrosome and kinetochore, and Plk4 lost its kinetochore function. The polo box arose N-terminal to the ancestral Plk kinase domain after Aurora split off. The closest relative of Mps1 was Tlk (bootstrap:36/100). The closest homolog of MadBub is an uncharacterized group of kinases. Interestingly, in contrast to their kinase domain, the TPR domains of Mps1 and Madbub are most closely related to one another, as indicated by the profile-versus-profile similarity searches (Table S2). This implied that the Mps1 and MadBub TPR domains joined with a kinase domain independently, as we observed before [35].

Zw10 homologs are involved in vesicle transport [36–38]. Its closest homolog is Cog5, which is involved in intra-Golgi transport (Figure S1A). Zw10 participates in two complexes: RZZ (Rod-Zwilch-Zw10), localized to the kinetochore, and the NRZ (Nag-Rint1-Zw10), involved in Golgi to ER transport. Notably, Rod is most closely related to Nag (Figure S1H), suggesting their ancestor interacted with Zw10 before it duplicated to give rise to Rod and Nag. Whether this ancestral complex was involved vesicle transport or in the kinetochore, or in both, is unclear.

The relatives of the WD40 kinetochore proteins are highly diverse, and their repetitive nature made it hard to resolve their (deep) evolutionary origins. Cdc20, a WD40 repeat protein, is most closely related to Cdh1 (Figure S1B), which like Cdc20 activates the APC/C [39]. Bub3’s closest homolog is Rae1 (Figure S1C), a protein involved in transporting mRNAs out of the nucleus [40]. For both Cdc20 and Bub3, we cannot suggest nor exclude that their ancestors were part of the kinetochore network. Regarding the deep origin of the WD40 repeat, it is not known yet if this domain already existed in prokaryotes before the pre-eukaryotic lineage, or if it was invented between FECA and LECA. While this repeat is clearly present in current-day prokaryotes [41], these may have received it recently from eukaryotes via horizontal gene transfer. TPR domains have been found in many prokaryotes and were suggested to have been present in the prokaryotic ancestors of eukaryotes [42].

#### Unique domains in the kinetochore?

Like the Mis12-like proteins, various other proteins domains such as Ska seem unique to the kinetochore (Table 1). While these domains might have originated between FECA and LECA and only serve roles in the kinetochore, we cannot exclude that they do have homologous prokaryotic or eukaryotic sequences, but that we are not able to detect these. The same possibility applies to those kinetochore proteins for which we do not have indications for any homologs at all, such as Zwilch, Incenp, Borealin, Shugoshin, Cep57, CenpH, CenpK, CenpQ and CenpU.

### Mosaic origin of the LECA kinetochore

Most LECA kinetochore proteins consisted of domains found in other eukaryotic proteins (37/55, 67%), while others had no detectable homology outside of the kinetochore (18/55, 33%, Table 1, Figure 1B). From the proteins with common domains, only one (Trip13) was directly derived from its prokaryotic ancestors. All others have eukaryotic homologs (paralogs) that are more closely related than prokaryotic homologs (if any). These paralogs are involved in an array of eukaryotic cellular processes (Table 1, last two columns). Altogether, the ancient homologs of kinetochore proteins indicate that the kinetochore has a mosaic origin. Specific eukaryotic processes were prevalent amongst the evolutionary links. Of the 15 closest non-kinetochore homologs that we identified (Table 1, Figure 5), five were involved in chromatin and/or transcription regulation (Tlk1, H3, Rev7 Med15, FancL), two played a role in Golgi and ER-related vesicle transport systems (Nag, Cog5) and another two are associated with centriole biogenesis (Cluap1, Plk4). More distantly related homologs were involved in DNA repair and replication (Dpoe3-4 and the replication factors: Cdt1, Cdc6 and Orc1), chromatin structure (nucleosomal histones), transcriptional regulation (e.g. TBP, TAFs, CBF/NF, NC2), RNA splicing (Fam98, Syf1/Crooked neck-like), vesicle transport (Kif1C, AP-2/4B, CopG1, AP-1G, CopB, Rab1A, Ccdc22, Ccdc93) and intra-flagellar transport (Ift54, Ift81). All in all, most LECA kinetochore proteins are part of families that have many members in eukaryotes, like UBCs, kinases and histones. Such families dramatically expanded between FECA and LECA and diversified into different eukaryotic cellular processes, including the kinetochore.

**Figure 5.**
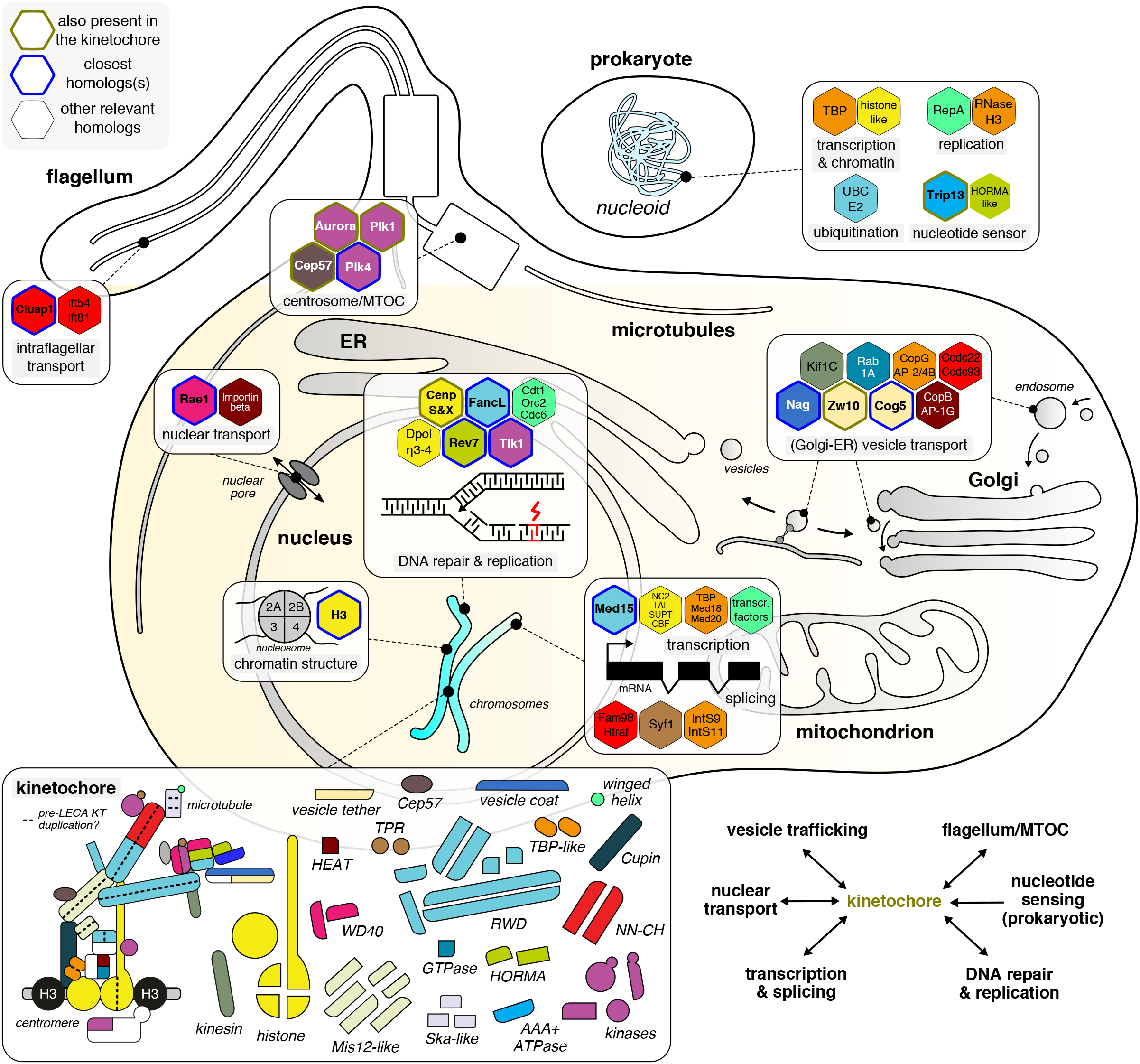
Mosaic origin of the eukaryotic kinetochore. Overview of the eukaryotic and prokaryotic (closest) homologs of LECA kinetochore proteins, which play a role in a great variety of cellular processes, signifying the mosaic origin of the eukaryotic kinetochore. Relevant eukaryotic and prokaryotic homologs (hexagons) of LECA kinetochore proteins are coloured based on the presence of a shared domain (see overview of the parts at the bottom), and projected onto the location(s) in the eukaryotic cell at which they operate (see for detailed information Table 1). To indicate the homologous and/or functional relationship with kinetochore proteins, the hexagons of homologs are lined with different colours to indicate: (bold green) a LECA kinetochore protein that also has a non-kinetochore function, (bold blue) the closest homolog to a LECA kinetochore protein and (thin black) more distantly related homologs of LECA kinetochore proteins. In addition, distantly related homologs of TBP-like, histones, UBC/RWD and HORMA domain-containing kinetochore proteins were already present in prokaryotes (top right). Bottom (left): overview of the different number and types of domains in the LECA kinetochore. The dotted lines indicate a potential intra-kinetochore duplication during eukaryogenesis leading to the formation of various heteromeric (sub)complexes within the kinetochore. (bottom right) summary of the evolutionary links between prokaryotic/eukaryotic molecular systems and the kinetochore.

In addition to their mosaic origins, many kinetochore proteins arose from intra-kinetochore gene duplications. Of the 40 kinetochore domains with an identified closest homolog (as referred to in ‘Identifying ancient homologs of kinetochore proteins’), 27 (68%) are most closely related to another kinetochore protein, indicating an important role for intra-kinetochore duplications in its evolutionary origin (Table 1). We inferred that the 55 domains result from 36 ancestral kinetochore units (‘anc_KT’ units), implying that intra-kinetochore gene duplications expanded the primordial kinetochore by a factor of ~1.5. We observed few domain fusions among LECA KT proteins. In fact, we find three: in Mps1 and MadBub, whose TPR domains independently joined their kinase domains, and a fusion of a microtubule-binding winged-helix and a Ska-like domain in Ska1 (see Table S3).

## Discussion

### Evolution of eukaryotic cellular systems

We have here shown that the kinetochore largely consists of paralogous proteins, which either share deep evolutionary roots with a variety of other core eukaryotic cellular processes or are novel and specific to the kinetochore itself (Mis12 and Ska) (Figure 5). In the origin of the kinetochore, gene duplications played a key role, which is in line with a previously reported elevated rate of gene duplications in eukaryogenesis [43]. Duplications contributed to the expansions of e.g. the spliceosome [12], the intraflagellar transport complex [44], COPII [45] and the nuclear pore [14]. However, the role of duplications in the origin ofthe kinetochore is different from their role in membrane-specifying complexes, in which paralogs are mainly shared between the different organelles rather than within them [46]. In tethering complexes, duplications generated proteins both within and between complexes [36]. Kinetochore proteins with prokaryotic ancestry sometimes conserved certain prokaryotic biochemical functions (e.g. HORMA-Trip13 interaction, histone-DNA interaction by CenpA) but no longer perform the ancestral cellular function. Therefore, the kinetochore followed a different evolutionary trajectory between FECA and LECA than e.g. NADH:ubiquinone oxidoreductase (Complex I) [47], which was directly derived from the Alphaproteobacterium that became the mitochondrion (Figure 1), and expanded between FECA and LECA by incorporating additional proteins of different origins. The Golgi and ER also differ from the kinetochore, as they mainly have archaeal roots [48]. The nuclear pore, while resembling the kinetochore in having a mosaic origin, was assembled with a substantial number of proteins derived from prokaryotic sequences [12, 14]. The latter is also true for the spliceosome [12, 14].

### Intra-kinetochore duplication

The intra-kinetochore duplications suggest an evolutionary trajectory by which the kinetochore partially expanded through homodimers that became heterodimers via gene duplication [49]. A primordial kinetochore might have been composed of complexes that consisted of multimers of single ancestral proteins (‘anc_KT’ in Table 1). After these proteins duplicated, the resulting paralogs maintained the capacity to interact, resulting in a heteromer. For example, the Ndc80 complex might have consisted of a tetramer of two copies of an ancient CH protein and two copies of an ancient RWD protein. According to this model, the proteins with shared domains within complexes should be most closely related to one another. This paradigm holds for the Ska subunits, the CH domain proteins, TBP-like proteins and the RWD proteins, and partially for the Mis12-like proteins (those within the Mis12 complex) and the histone fold proteins (CenpS-X:CenpT-W). We observe that many paralogous proteins are positioned along the inner-outer kinetochore axis (Figure 5, dashed line). We speculate that not too long before LECA, the genes encoding the proteins along this axis duplicated in quick stepwise succession or in one event [49–51], which would be consistent with the proposed syncytial nature of lineages that gave rise to LECA [52].

### Rapid sequence evolution of kinetochore components

The LECA kinetochore contains protein domains that are unique to the kinetochore and therefore, by definition, unique to eukaryotes (33% of LECA kinetochore protein domains). New and more diverse genomes or elucidated protein structures may allow for the detection of such distant homologs in the future. Kinetochore proteins that do share domains with other eukaryotic systems, such as the RWD, TBP-like, histones and TPR, seem to be strongly diverged in the kinetochore. For example, the TPR domains of Mps1 and MadBub are more derived than those of the Anaphase Promoting Complex/Cyclosome (APC/C). This suggests that, after these domains got involved in the kinetochore, their sequences evolved more rapidly, and continued to do so after LECA [6]. Rapid evolution after LECA may be correlated with the widespread rapid divergence of centromere sequences. An evolutionary acceleration may also have occurred to the ‘*de novo*’ proteins in the LECA kinetochore, causing homology detection to fail.

### Possible origins of the kinetochore during eukaryogenesis

Tracing in what order these proteins or domains got involved in the kinetochore, relative to the origin of other eukaryotic features, would be highly interesting. Possibly, an early, very basic kinetochore was just composed of the centromere- and microtubule-binding proteins, similar to prokaryotic systems, while the CCAN (the ‘Cenp’ proteins), which serves as their bridge, was added later. Relative timings of such attributions could potentially shed light on the evolution of eukaryotic chromosome segregation. Although little is known about evolution of the eukaryotic segregation machinery, it must be associated to the evolution of linear chromosomes, the evolution of the nucleus and of the eukaryotic cytoskeleton, including centrosomes. Because the kinetochore shares ancestry with many other eukaryotic processes and cellular features and therefore does not seem to have an explicit prokaryotic or eukaryote template structure or process, we envision it originated late during eukaryogenesis. The evolutionary link with flagellar transport systems, may signify an early role for the flagellum in coordinating microtubule-based mechanisms of chromosome segregation, which is consistent with the function of the centriole as the microtubule organizing centre in most eukaryotes. A common origin with Golgi/ER-related vesicle transport components could potentially point to membrane-based mechanisms of chromosome segregation in pre-LECA lineages, similar to those found in prokaryotes (Figure 1A). Because currently no eukaryotes or ‘proto’-eukaryotes are known that might segregate chromosomes in a pre-LECA manner, it remains hard to unravel which series of events gave rise to the spindle apparatus, the centromere and the kinetochore. The currently known closest archaeal relatives of eukaryotes, the Asgard Archaea [53, 54] (Figure 1A), clearly do not operate a eukaryote-like chromosome segregation system, but unidentified closer related prokaryotes or proto-eukaryotes could. New (meta)genomic sequences aided in reconstructing the evolution of the ubiquitin system [55] and the membrane trafficking system [48]. Similarly, such newly identified species may enhance our understanding of the pre-LECA evolution of the eukaryotic kinetochore and the chromosome segregation machinery.

## Data and Methods

See Supplementary Data and Methods

## Author contributions

JJEH and ET performed the research. JJEH, ET, BS and GK conceived the project and wrote the manuscript.

## Acknowledgments

We thank Leny van Wijk for providing the phylogenetic tree of eukaryotic kinases and for helping to construct the eukaryotic proteome database, for which we also thank John van Dam. We thank the members of the Kops and Snel labs for helpful discussions on the research. This work was supported by Netherlands Organisation for Scientific Research (NWO-Vici 016.160.638 to BS). ET is supported by a postdoctoral fellowship from the Herchel Smith Fund, Cambridge, UK.

## Supplementary Information

Supplementary Information consists of Supplementary Data and Methods, Supplementary Text, Supplementary Figures (4), Supplementary Tables (4) and Supplementary Files (154).

## References

1. Makarova M, Oliferenko S (2016) Mixing and matching nuclear envelope remodeling and spindle assembly strategies in the evolution of mitosis. Curr Opin Cell Biol 41:43–50. doi: 10.1016/j.ceb.2016.03.016

2. De Souza CPC, Osmani SA (2007) Mitosis, Not Just Open or Closed. Eukaryot Cell 6:1521–1527. doi: 10.1128/ec.00178-07

3. Drechsler H, McAinsh AD (2012) Exotic mitotic mechanisms. Open Biol 2:120140. doi: 10.1098/rsob.120140

4. Sazer S, Lynch M, Needleman D (2014) Deciphering the Evolutionary History of Open and Closed Mitosis. Curr Biol 24:1099–1103. doi: 10.1016/j.cub.2014.10.011

5. Henikoff S, Ahmad K, Malik HS (2001) The centromere paradox: stable inheritance with rapidly evolving DNA. Science 293:1098–1102. doi: 10.1126/science.1062939

6. van Hooff JJ, Tromer E, van Wijk LM, et al (2017) Evolutionary dynamics of the kinetochore network in eukaryotes as revealed by comparative genomics. EMBO Rep 18:1559–1571. doi: 10.15252/embr.201744102

7. Barillà D (2016) Driving Apart and Segregating Genomes in Archaea. Trends Microbiol 24:957–967. doi: https://doi.org/10.1016/j.tim.2016.07.001

8. Badrinarayanan A, Le TBK, Laub MT (2015) Bacterial Chromosome Organization and Segregation. Annu Rev Cell Dev Biol 31:171–199. doi: 10.1146/annurev-cellbio-100814-125211

9. Lindås A-C, Bernander R (2013) The cell cycle of archaea. Nat Rev Microbiol 11:627–638. doi: 10.1038/nrmicro3077

10. Dacks JB, Field MC, Buick R, et al (2016) The changing view of eukaryogenesis - fossils, cells, lineages and how they all come together. J Cell Sci 129:3695–3703. doi: 10.1242/jcs.178566

11. Koonin E V (2010) The origin and early evolution of eukaryotes in the light of phylogenomics. Genome Biol 11:209. doi: 10.1186/gb-2010-11-5-209

12. Vosseberg J, Snel B (2017) Domestication of self-splicing introns during eukaryogenesis: the rise of the complex spliceosomal machinery. Biol Direct 12:30. doi: 10.1186/s13062-017-0201-6

13. Field MC, Dacks JB (2009) First and last ancestors: reconstructing evolution of the endomembrane system with ESCRTs, vesicle coat proteins, and nuclear pore complexes. Curr Opin Cell Biol 21:4–13. doi: 10.1016/j.ceb.2008.12.004

14. Mans BJ, Anantharaman V, Aravind L, Koonin E V (2004) Comparative genomics, evolution and origins of the nuclear envelope and nuclear pore complex. Cell Cycle 3:1612–1637. doi: 1316 [pii]

15. Schmitzberger F, Harrison SC (2012) RWD domain: a recurring module in kinetochore architecture shown by a Ctf19-Mcm21 complex structure. EMBO Rep 13:216–222. doi: 10.1038/embor.2012.1

16. Doerks T, Copley RR, Schultz J, et al (2002) Systematic identification of novel protein domain families associated with nuclear functions. Genome Res 12:47–56. doi: 10.1101/

17. Burroughs AM, Jaffee M, Iyer LM, Aravind L (2008) Anatomy of the E2 ligase fold: implications for enzymology and evolution of ubiquitin/Ub-like protein conjugation. J Struct Biol 162:205–218. doi: 10.1016/j.jsb.2007.12.006

18. Petrovic A, Mosalaganti S, Keller J, et al (2014) Modular Assembly of RWD Domains on the Mis12 Complex Underlies Outer Kinetochore Organization. Mol Cell 53:591–605. doi: 10.1016/j.molcel.2014.01.019

19. Mattiroli F, Bhattacharyya S, Dyer PN, et al (2017) Structure of histone-based chromatin in Archaea. Science 357:609–612. doi: 10.1126/science.aaj1849

20. Malik HS, Henikoff S (2003) Phylogenomics of the nucleosome. Nat Struct Mol Biol 10:882–891

21. Zhao Q, Saro D, Sachpatzidis A, et al (2014) The MHF complex senses branched DNA by binding a pair of crossover DNA duplexes. Nat Commun 5:2987. doi: 10.1038/ncomms3987

22. Tao Y, Jin C, Li X, et al (2012) The structure of the FANCM-MHF complex reveals physical features for functional assembly. Nat Commun 3:782. doi: 10.1038/ncomms1779

23. Pentakota S, Zhou K, Smith C, et al (2017) Decoding the centromeric nucleosome through CENP-N. Elife 6:. doi: 10.7554/eLife.33442

24. Chittori S, Hong J, Saunders H, et al (2018) Structural mechanisms of centromeric nucleosome recognition by the kinetochore protein CENP-N. Science 359:339–343. doi: 10.1126/science.aar2781

25. Hinshaw SM, Harrison SC (2013) An Iml3-Chl4 heterodimer links the core centromere to factors required for accurate chromosome segregation. Cell Rep 5:29–36. doi: 10.1016/j.celrep.2013.08.036

26. Brindefalk B, Dessailly BH, Yeats C, et al (2013) Evolutionary history of the TBP-domain superfamily. Nucleic Acids Res 41:2832–2845. doi: 10.1093/nar/gkt045

27. Koster MJE, Snel B, Timmers HTM (2015) Genesis of Chromatin and Transcription Dynamics in the Origin of Species. Cell 161:724–736. doi: http://dx.doi.org/10.1016/j.cell.2015.04.033

28. Burroughs AM, Zhang D, Schäffer DE, et al (2015) Comparative genomic analyses reveal a vast, novel network of nucleotide-centric systems in biological conflicts, immunity and signaling. Nucleic Acids Res 43:10633–10654. doi: 10.1093/nar/gkv1267

29. Gimona M, Djinovic-Carugo K, Kranewitter Wolfgang J, Winder Steven J (2001) Functional plasticity of CH domains. FEBS Lett 513:98–106. doi: 10.1016/S0014-5793(01)03240-9

30. Schou KB, Andersen JS, Pedersen LB (2014) A divergent calponin homology (NN-CH) domain defines a novel family: Implications for evolution of ciliary IFT complex B proteins. Bioinformatics 30:899–902. doi: 10.1093/bioinformatics/btt661

31. Pasek RC, Berbari NF, Lewis WR, et al (2012) Mammalian Clusterin associated protein 1 is an evolutionarily conserved protein required for ciliogenesis. Cilia 1:1–20. doi: 10.1186/2046-2530-1-20

32. Pérez-González A, Pazo A, Navajas R, et al (2014) hCLE/C14orf166 associates with DDX1-HSPC117-FAM98B in a novel transcription-dependent shuttling RNA-transporting complex. PLoS One 9:e90957. doi: 10.1371/journal.pone.0090957

33. Healy MD, Hospenthal MK, Hall RJ, et al (2018) Structural insights into the architecture and membrane interactions of the conserved COMMD proteins. Elife 7:. doi: 10.7554/eLife.35898

34. Mallam AL, Marcotte EM (2017) Systems-wide Studies Uncover Commander, a Multiprotein Complex Essential to Human Development. Cell Syst. 4:483–494

35. Nijenhuis W, von Castelmur E, Littler D, et al (2013) A TPR domain-containing N-terminal module of MPS1 is required for its kinetochore localization by Aurora B. J Cell Biol 201:217–231. doi: 10.1083/jcb.201210033

36. Koumandou VL, Dacks JB, Coulson RMR, Field MC (2007) Control systems for membrane fusion in the ancestral eukaryote; evolution of tethering complexes and SM proteins. BMC Evol Biol 7:1–29. doi: 10.1186/1471-2148-7-29

37. Hong W, Lev S (2014) Tethering the assembly of SNARE complexes. Trends Cell Biol 24:35–43. doi: 10.1016/j.tcb.2013.09.006

38. Schroeter S, Beckmann S, Schmitt HD (2016) Coat/Tether Interactions-Exception or Rule? Front cell Dev Biol 4:1–44. doi: 10.3389/fcell.2016.00044

39. Pfleger CM, Lee E, Kirschner MW (2001) Substrate recognition by the Cdc20 and Cdh1 components of the anaphase-promoting complex. Genes Dev 15:2396–2407. doi: 10.1101/gad.918201

40. Murphy R, Watkins JL, Wente SR (1996) GLE2, a Saccharomyces cerevisiae homologue of the Schizosaccharomyces pombe export factor RAE1, is required for nuclear pore complex structure and function. Mol Biol Cell 7:1921–37

41. Hu XJ, Li T, Wang Y, et al (2017) Prokaryotic and Highly-Repetitive WD40 Proteins: A Systematic Study. Sci Rep 7:1–13. doi: 10.1038/s41598-017-11115-1

42. Schlegel T, Mirus O, von Haeseler A, Schleiff E (2007) The Tetratricopeptide Repeats of Receptors Involved in Protein Translocation across Membranes. Mol Biol Evol 24:2763–2774. doi: 10.1093/molbev/msm211

43. Makarova KS, Wolf YI, Mekhedov SL, et al (2005) Ancestral paralogs and pseudoparalogs and their role in the emergence of the eukaryotic cell. Nucleic Acids Res 33:4626–4638. doi: 10.1093/nar/gki775

44. van Dam TJP, Townsend MJ, Turk M, et al (2013) Evolution of modular intraflagellar transport from a coatomer-like progenitor. Proc Natl Acad Sci 110:6943–6948. doi: 10.1073/pnas.1221011110

45. Schlacht A, Dacks JB (2015) Unexpected Ancient Paralogs and an Evolutionary Model for the COPII Coat Complex. Genome Biol Evol 7:1098–1109. doi: 10.1093/gbe/evv045

46. Mast FD, Barlow LD, Rachubinski R a., Dacks JB (2014) Evolutionary mechanisms for establishing eukaryotic cellular complexity. Trends Cell Biol 24:435–442. doi: 10.1016/j.tcb.2014.02.003

47. Gabaldón T, Rainey D, Huynen MA (2005) Tracing the Evolution of a Large Protein Complex in the Eukaryotes, NADH:Ubiquinone Oxidoreductase (Complex I). J Mol Biol 348:857–870. doi: https://doi.org/10.1016/jjmb.2005.02.067

48. Klinger CM, Spang A, Dacks JB, Ettema TJG (2016) Tracing the Archaeal Origins of Eukaryotic Membrane-Trafficking System Building Blocks. Mol Biol Evol 33:1528–1541. doi: 10.1093/molbev/msw034

49. Pereira-Leal JB, Levy ED, Kamp C, Teichmann SA (2007) Evolution of protein complexes by duplication of homomeric interactions. Genome Biol 8:1–12. doi: 10.1186/gb-2007-8-4-r51

50. Dacks JB, Peden AA, Field MC (2009) Evolution of specificity in the eukaryotic endomembrane system. Int. J. Biochem. Cell Biol. 41:330–340

51. Dacks JB, Field MC (2018) Evolutionary origins and specialisation of membrane transport. Curr. Opin. Cell Biol. 53:70–76

52. Garg SG, Martin WF (2016) Mitochondria, the Cell Cycle, and the Origin of Sex via a Syncytial Eukaryote Common Ancestor. Genome Biol Evol 8:1950–1970. doi: 10.1093/gbe/evw136

53. Zaremba-Niedzwiedzka K, Caceres EF, Saw JH, et al (2017) Asgard archaea illuminate the origin of eukaryotic cellular complexity. Nature 541:353–358. doi: 10.1038/nature21031

54. Spang A, Saw JH, Jørgensen SL, et al (2015) Complex archaea that bridge the gap between prokaryotes and eukaryotes. Nature 521:173–179. doi: 10.1038/nature14447

55. Grau-Bove X, Sebe-Pedros A, Ruiz-Trillo I (2015) The eukaryotic ancestor had a complex ubiquitin signaling system of archaeal origin. Mol Biol Evol 32:726–739. doi: 10.1093/molbev/msu334

56. Petrovic A, Keller J, Liu Y, et al (2016) Structure of the MIS12 Complex and Molecular Basis of Its Interaction with CENP-C at Human Kinetochores. Cell 167:1028–1040. doi: 10.1016/j.cell.2016.10.005

57. Zhou X, Zheng F, Wang C, et al (2017) Phosphorylation of CENP-C by Aurora B facilitates kinetochore attachment error correction in mitosis. Proc Natl Acad Sci 114:10677–10676. doi: 10.1073/pnas.1710506114

