## Supplementary_Figures for "Mosaic origin of the eukaryotic kinetochore"

### Supplementary Figure 1A

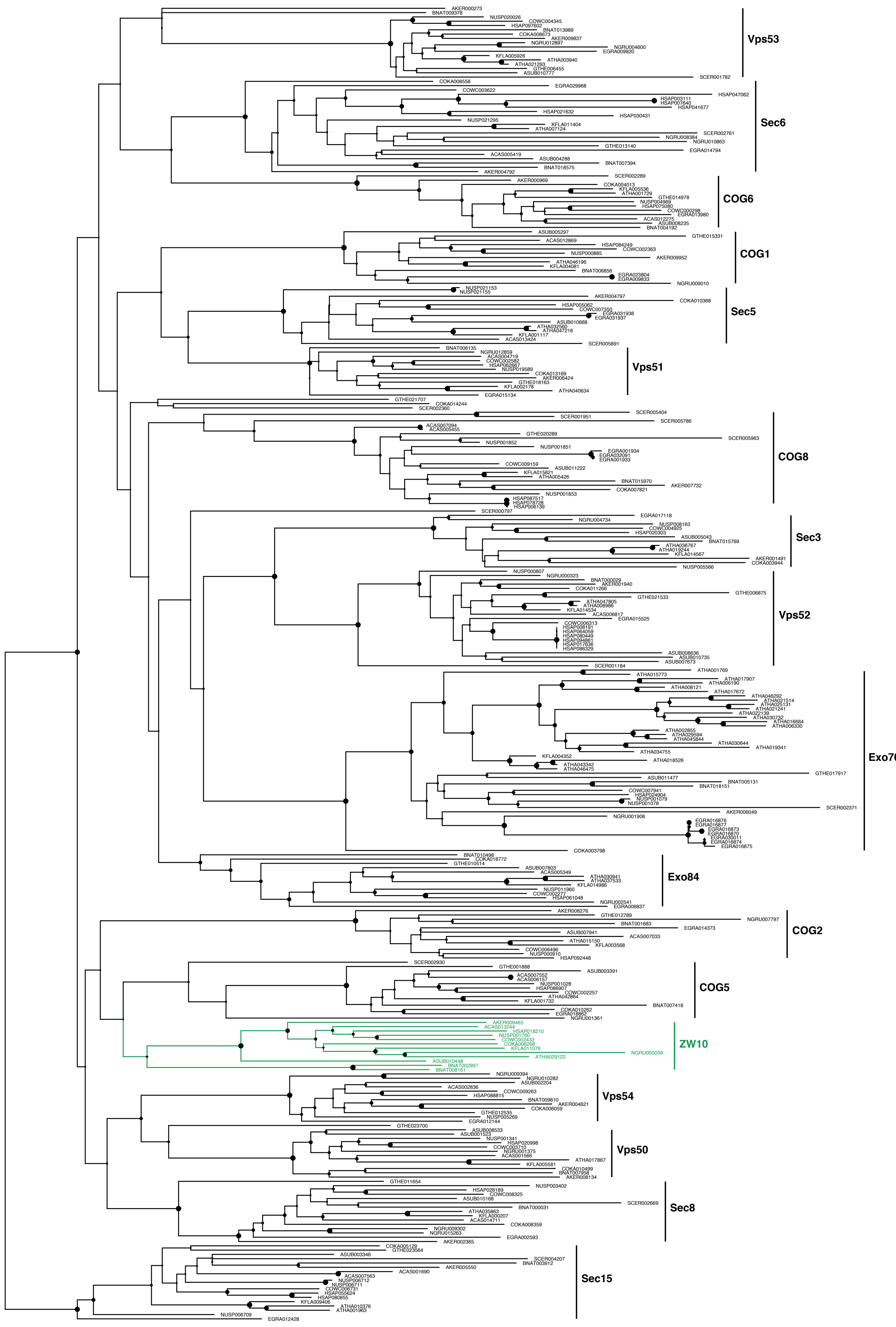


Support (rapid bootstraps)

0.6

Model: LG + G (RAxML)

### Supplementary Figure 1B

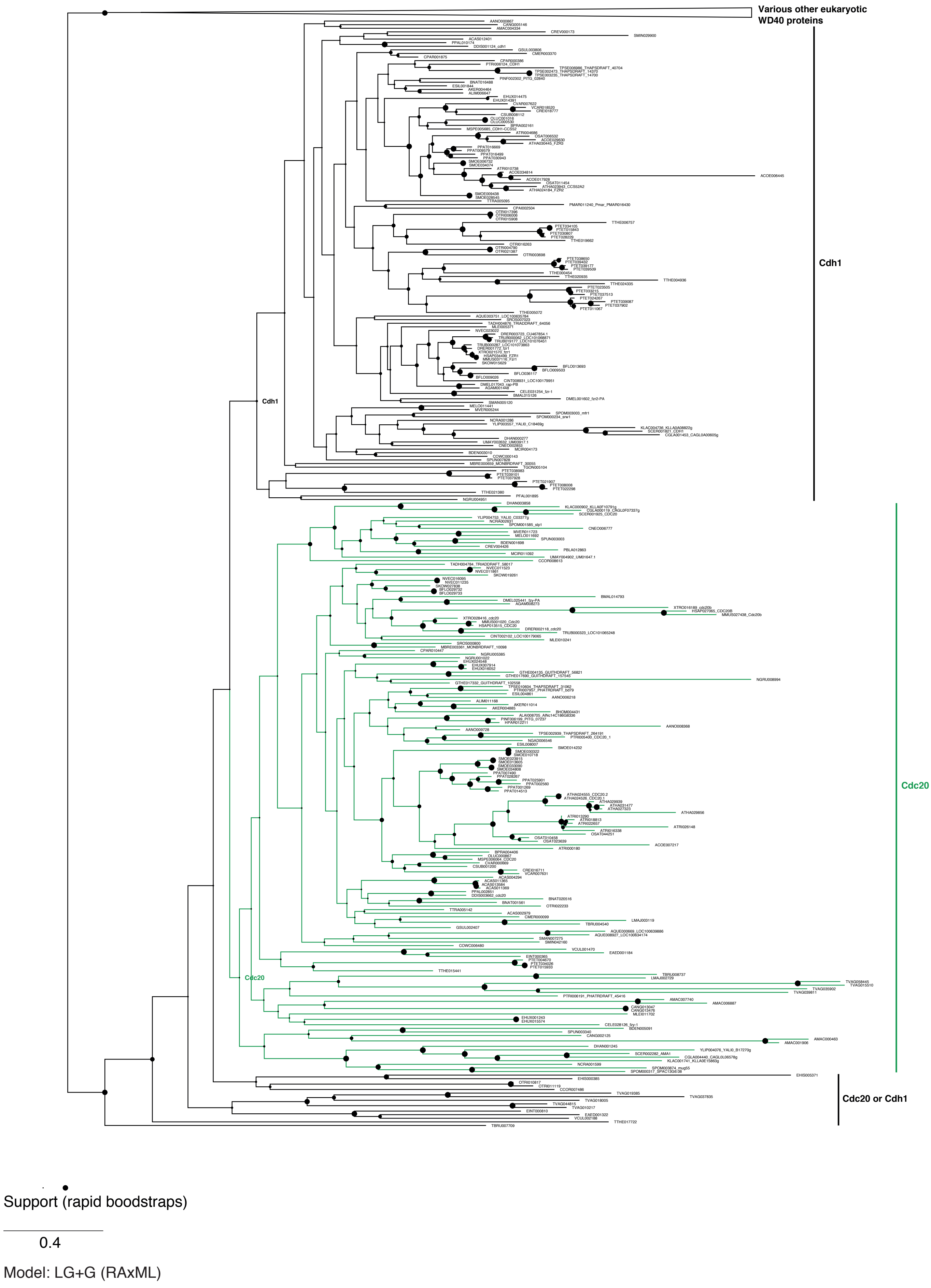

### Supplementary Figure 1C

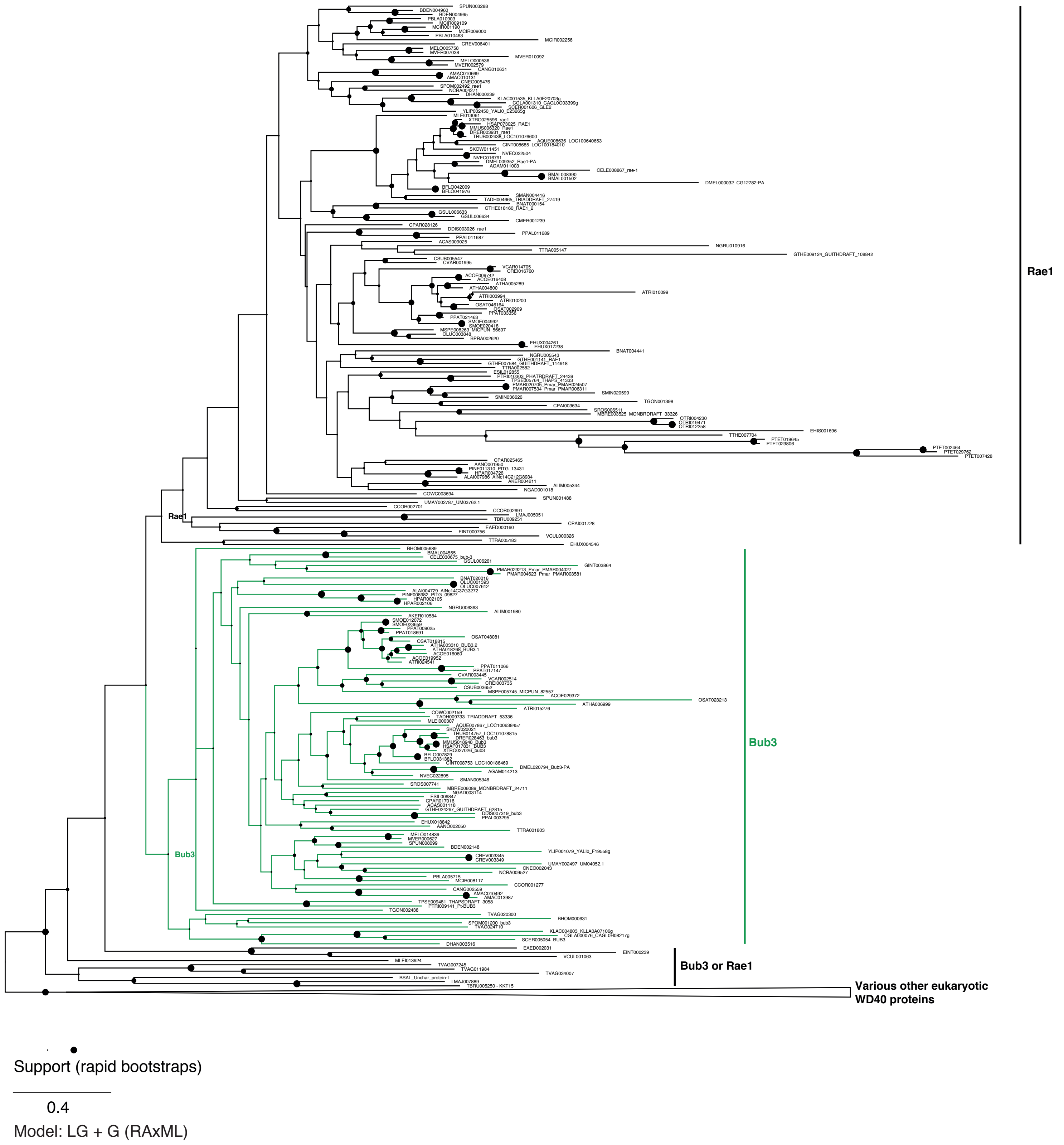

### Supplementary Figure 1D

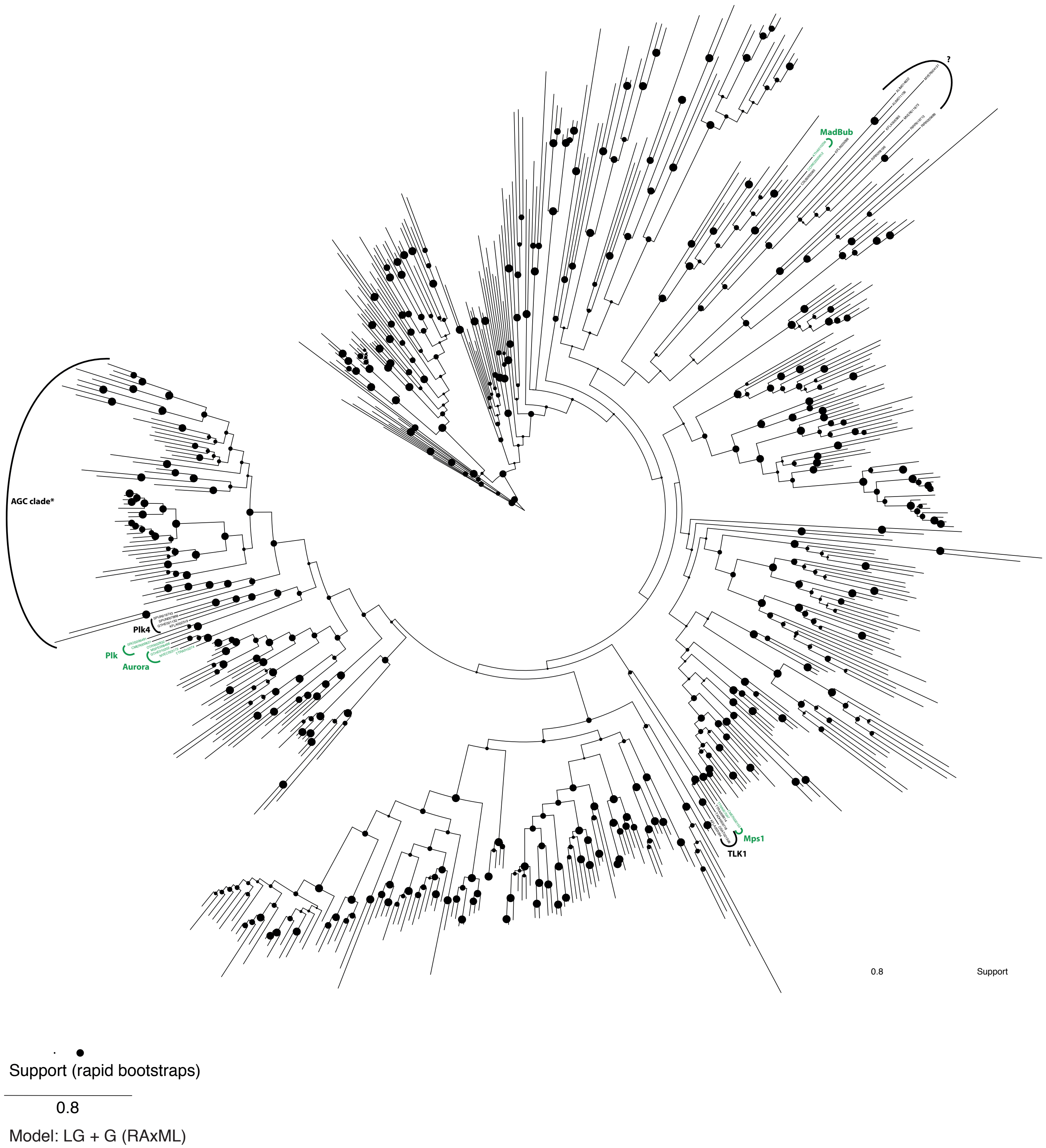

### Supplementary Figure 1E

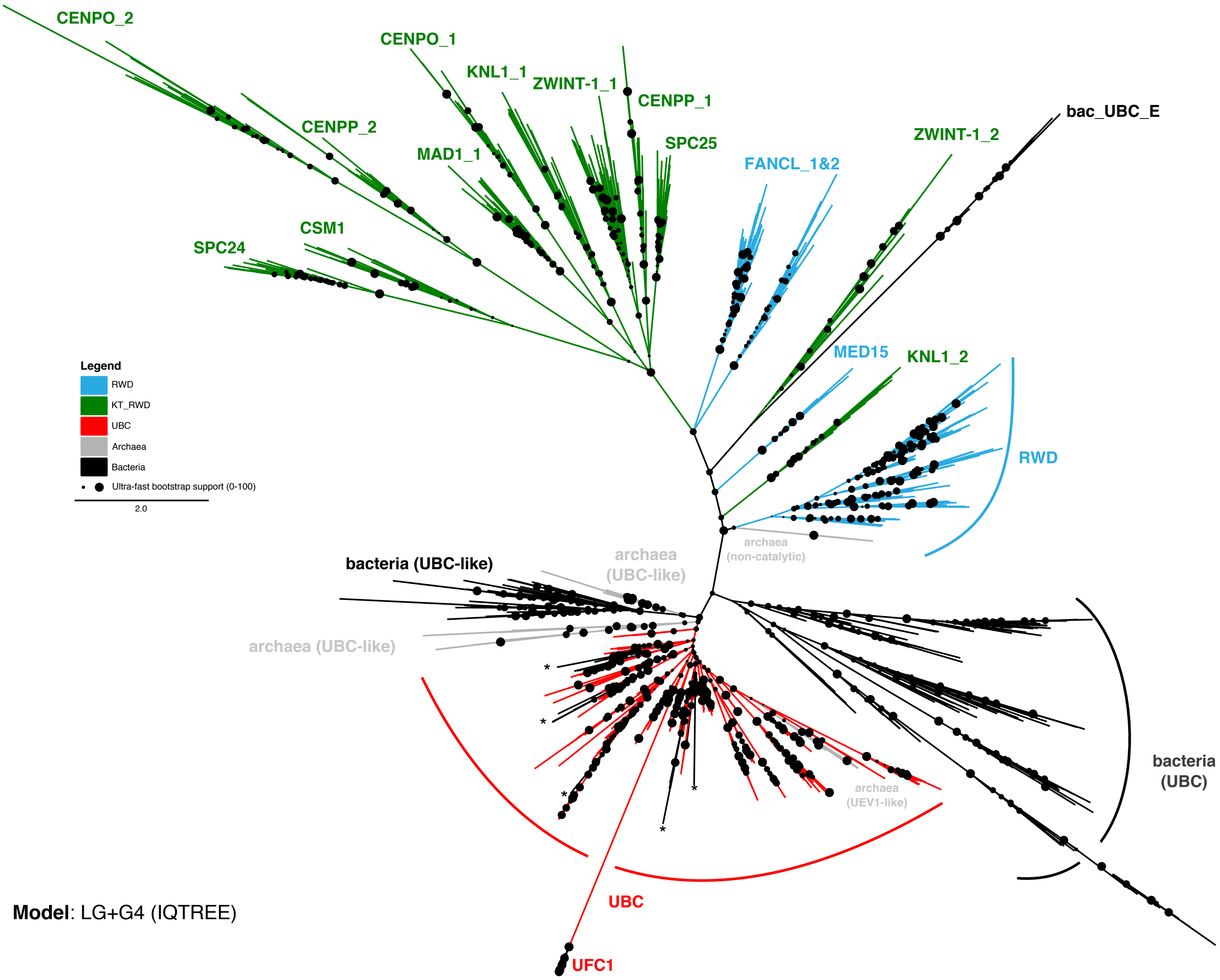


**Model**: LG+G4 (IQTREE)

### Supplementary Figure 1F

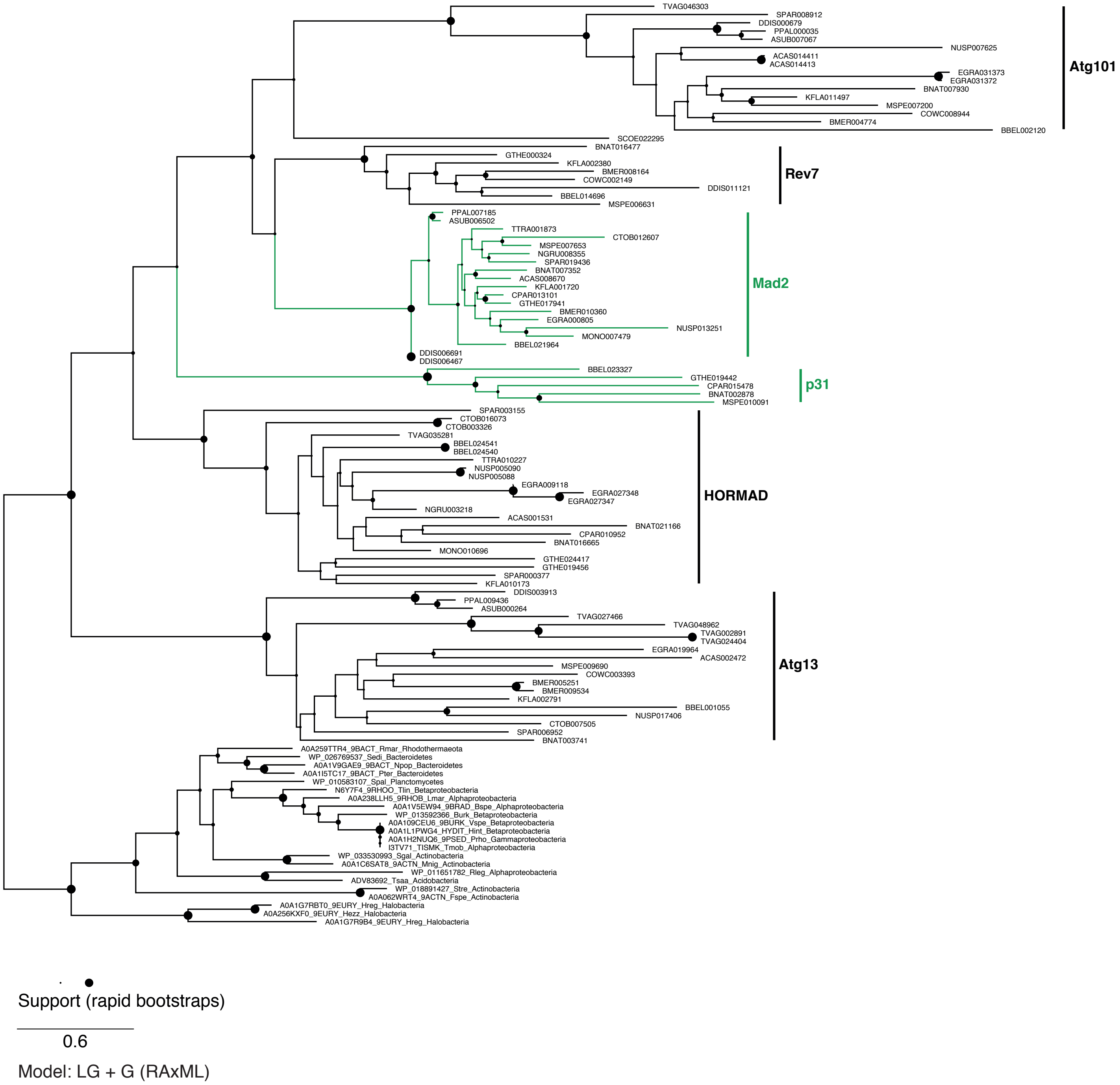

### Supplementary Figure 1G

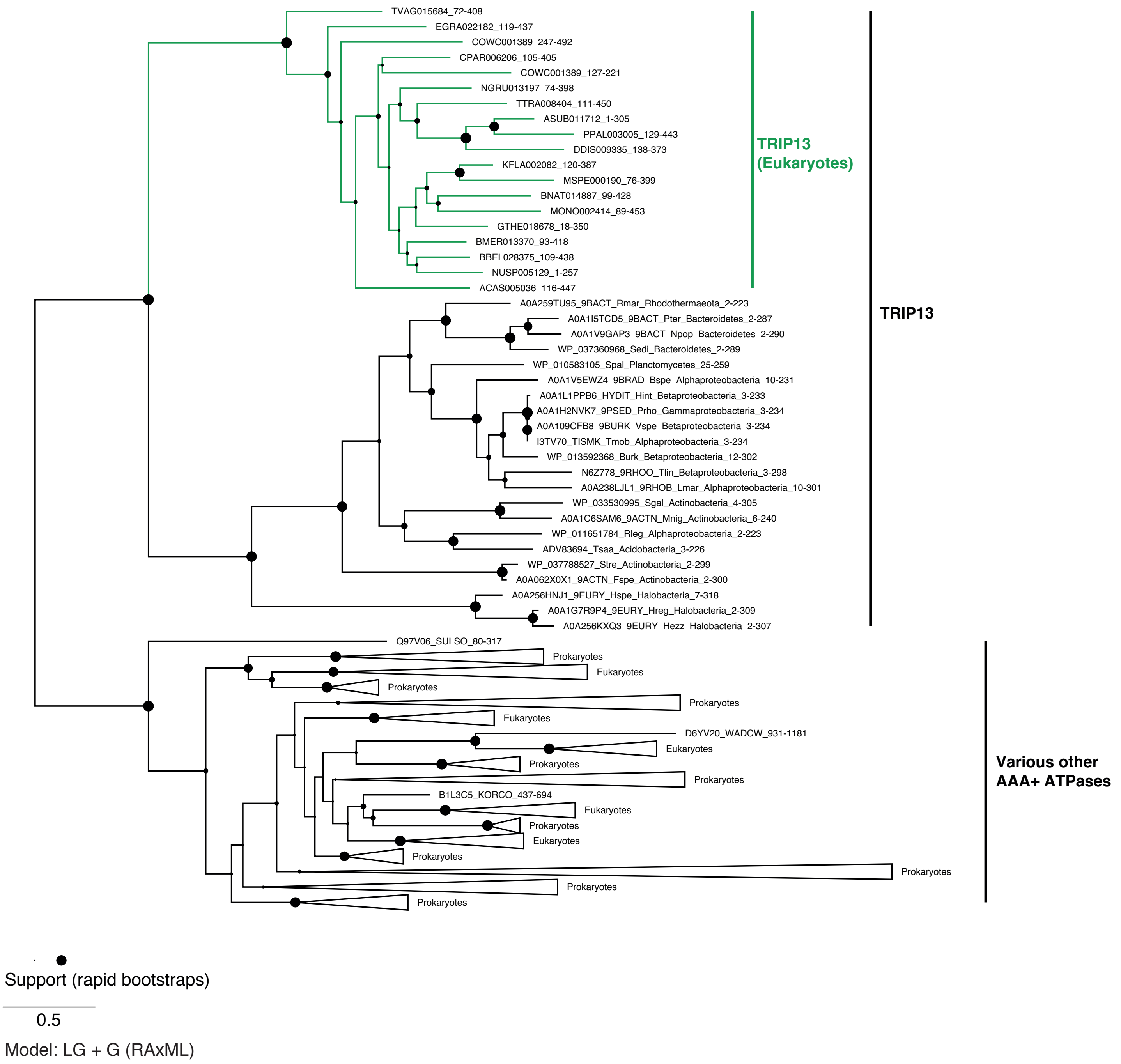

### Supplementary Figure 1H

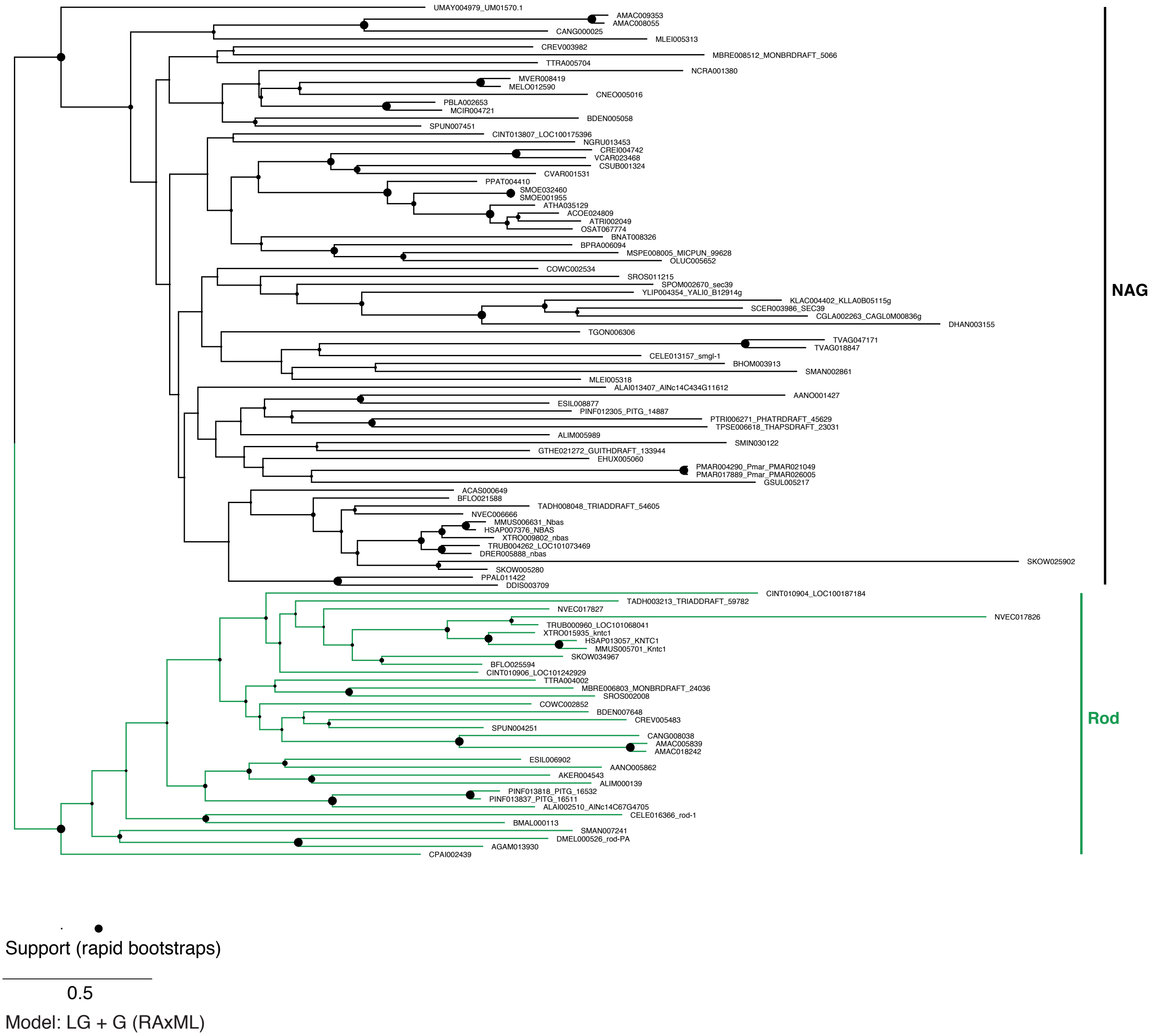

### Supplementary Figure 1I

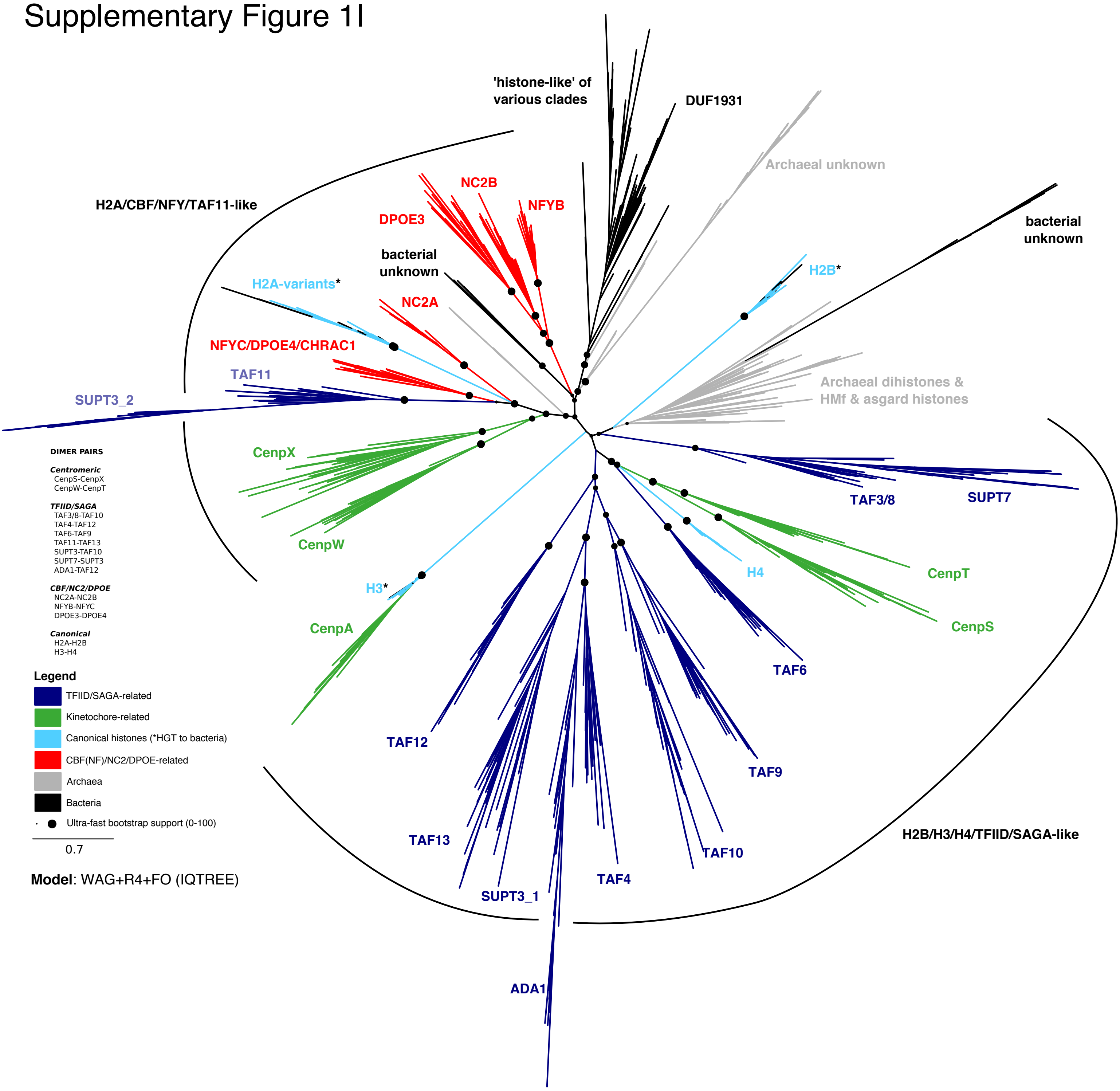


**Model**: WAG+R4+FO (IQTREE)

### Supplementary Figure 2

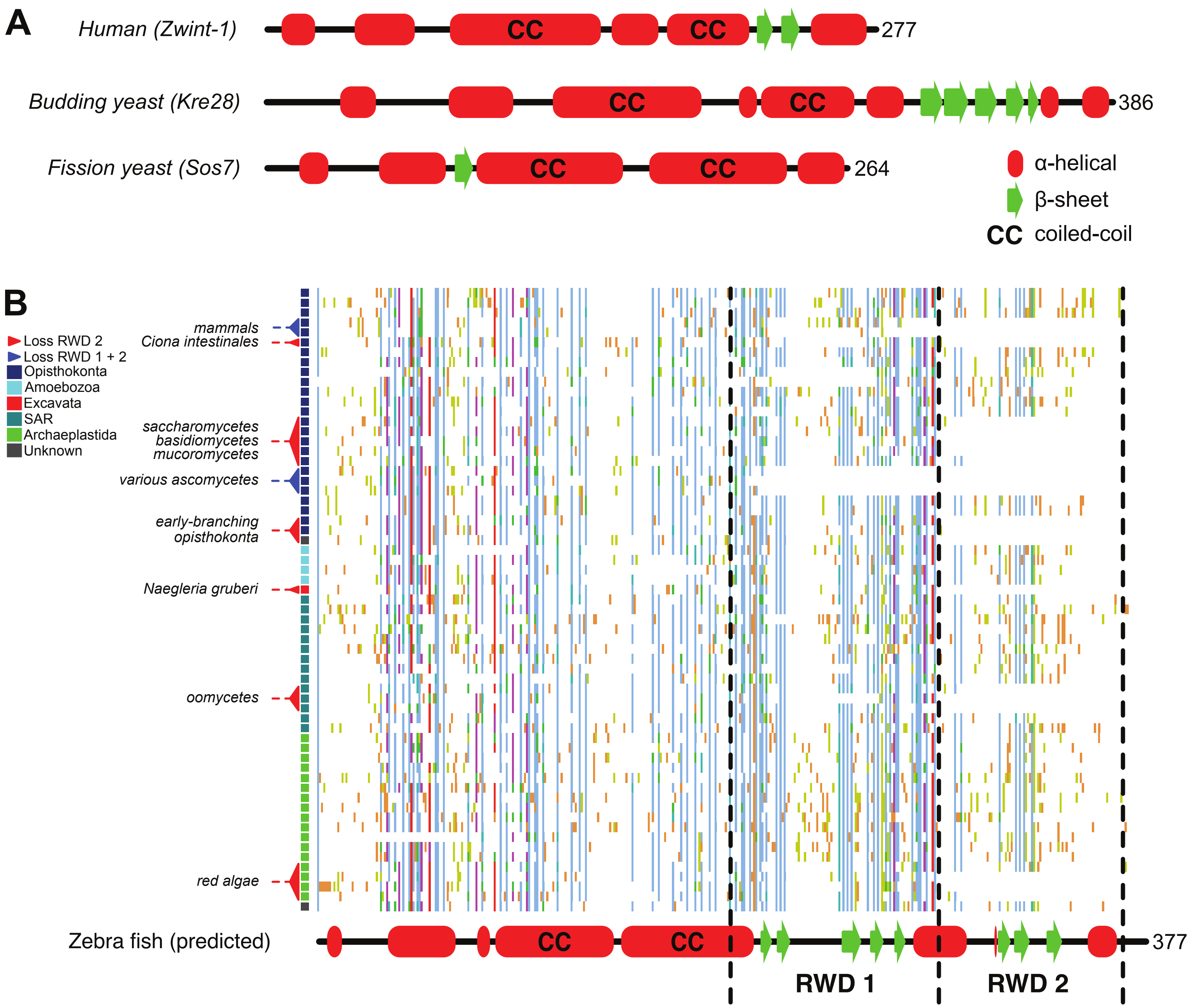

### Supplementary Figure 3

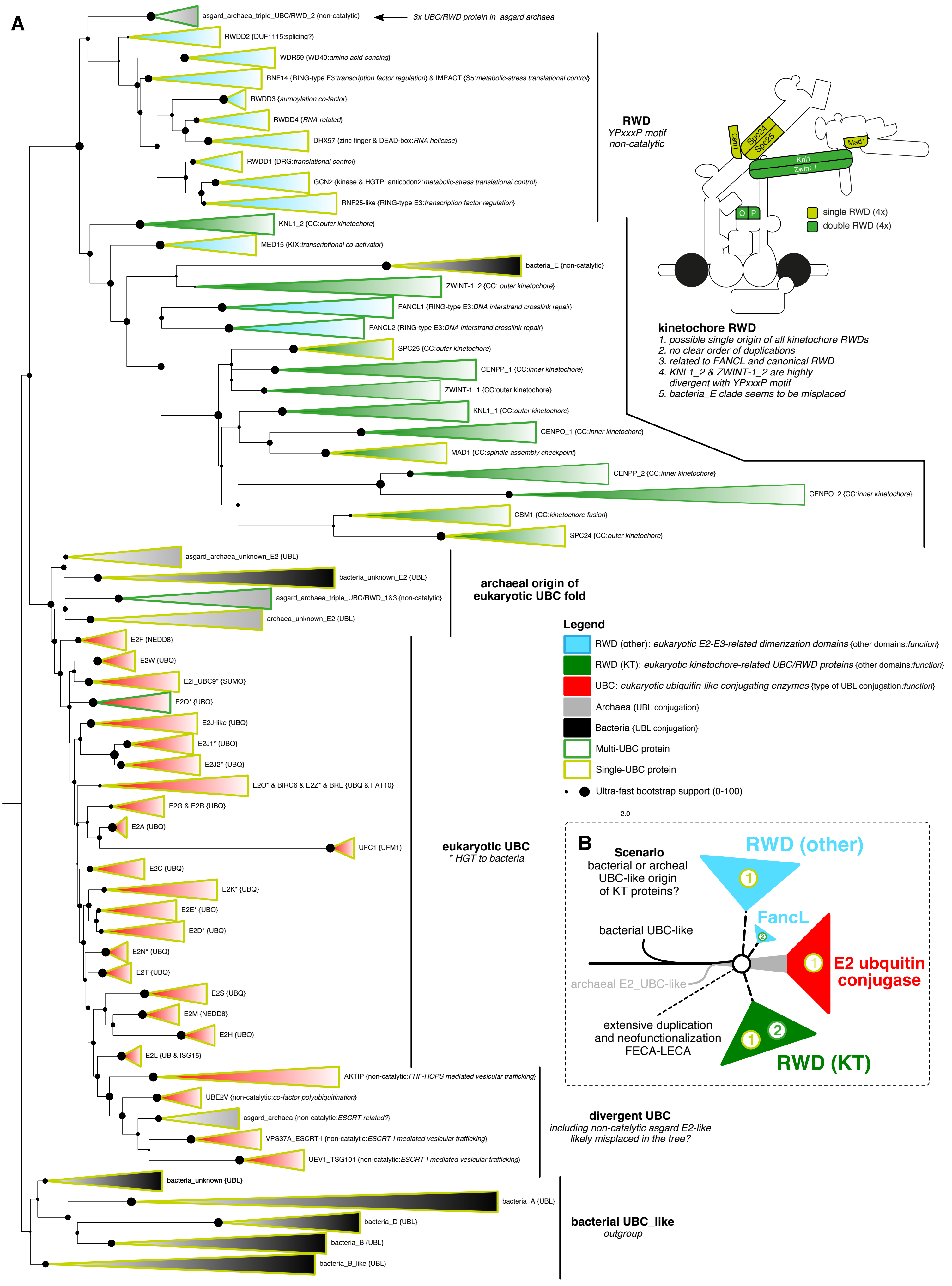

### Supplementary Figure 4

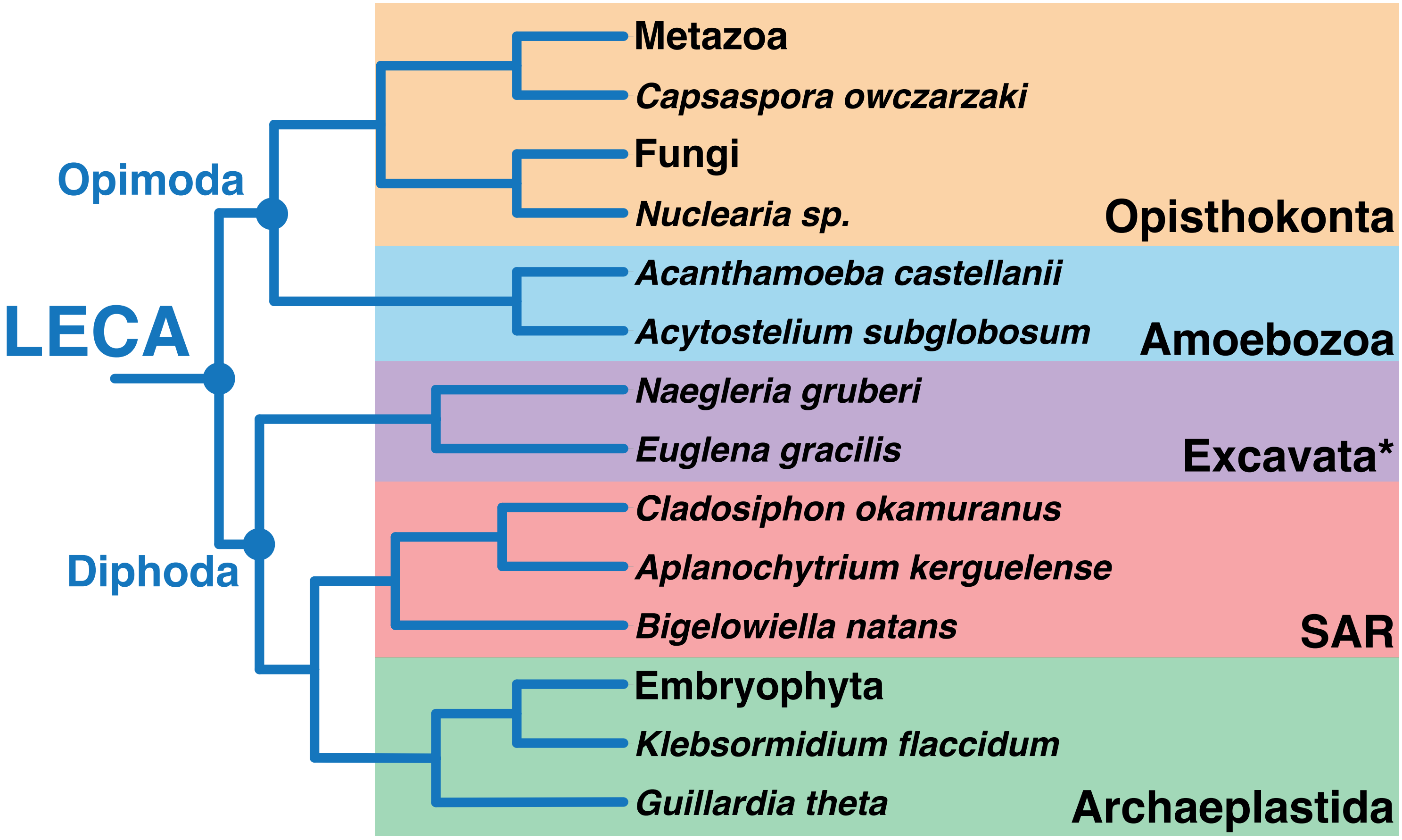
