## Supplementary_Information for "Mosaic origin of the eukaryotic kinetochore"

### Supplementary Data and Methods

#### Profile-versus-profile searches

To find distant homologs of kinetochore proteins, we constructed HMM profiles and used genome-wide databases to apply profile-versus-profile searches. For each of the proteins included in our previous analysis [6], and for some other proteins we studied more recently (Nkp1, Nkp2, Csm1, Lsr4, Mam1, Hrr25), we aligned the sets of orthologous sequences (MAFFT, v.7.149b [1] 'einsi' or 'linsi') and used these to construct HMM profiles (www.hmmer.org, version HMMER 3.1b1) and hhm-formatted profiles. For each protein, we made such a profile from the full-length alignment. In addition, if a protein has well-annotated domains, we made separate profiles of these domains. While Zwint-1 has two RWD domains, we only used the first as a separate domain profile, because the second is very poorly conserved across species (Figure S2). All HMM3 profiles can be found in Files S1-S147. We applied two different search strategies, using different tools and different search databases. For the first, we searched with full-length profiles of the kinetochore proteins and searched against a database compiled of profiles from PANTHER11.1 [2] and the kinetochore protein profiles themselves. For this search, we made use of of PRC (version 1.5.6) [3]. For the second strategy, we search with domain profiles if available for a given protein, and otherwise full-length profiles. We downloaded scop70 (March 1, 2016), pdb70 (September 14, 2016) and PfamA (version 31.0) profile databases from the HH-suite depository (ftp://toolkit.genzentrum.lmu.de/HH-suite/databases/hhsuite_dbs, downloaded on July 15, 2017) and combined these profiles with the kinetochore domain/full-length profiles we also used as queries. We searched using HHsearch (version 2.0.15) [4]. For each of the search strategies, we identified which of the database profiles correspond to the kinetochore domains/proteins. Using this information, we parsed the search results identifying 'best hits' and 'bidirectional best hits' for each kinetochore domain/protein profile. We made a network using te results from the second (HHsearch-based) search strategy, applying an E-value cut-off of 1 or 10. We visualised this network in Cytoscape (version 3.5.1) [5]. The network can be found in File S148. In this network, we also added the available cellular component GO terms to the hits using SIFTS [6]. In addition, we used information from the first (PRC-based) search strategy to trace the distant homology between subunits of the Mis12 and Nkp complexes (see also SI Text). The results of both search strategies can be found in Table S2.

#### Phylogenetic trees

For inference of the phylogenetic trees presented in this manuscript, we used a variety of methods. We collected homologs by searching with our tailor-made and Pfam HMM profiles against our local proteome database [7]. The first four letters of the eukaryotic sequences represent the species, of which the full names can be found in

Table S4. For the phylogenies of Zw10 and related tethering factors, HORMA, histones, RWD and Trip13, we used a subset of the species in this database, including species present in Figure S4. The phylogeny of all eukaryotic kinases was based on sequence-based subsampling, selecting for the slowest-evolving kinases [8]. For the prokaryotic sequences in the UBC/RWD and histone phylogenies, we performed phmmer/jackhmmer online and collected sequence hits from the uniprot database. For the HORMA and Trip13 phylogenies, we searched with the bacterial sequences reported by Burroughs et al. [9], from species possessing 'Bacterial_HORMA2' (for sequences annotated as 'Bacterial_HORMA1' we were not able to detect sufficient sequence similarity to construct a good-quality multiple sequence alignment, therefore we excluded these Bacterial_HORMA1 sequences as well as the Trip13 proteins in the respective species). We added contig information for species not included by Burroughs et al. [9] (see File S154 for the HORMA-Trip13 operon that we found for *Haloarchaea*, *Rhodothermaeotum* and *Acidobacteria*). For all protein families, multiple sequence alignments were inferred using MAFFT (v.7.149b [1], 'einsi' or 'linsi') [10], and trimmed with trimAl (1.2rev59, various options) [11]. Due to the high degree of sequence divergence for the UBC and histone fold-containing proteins, we constructed a super alignment of trusted trimmed orthologous group alignments using the function 'merge' of MAFFT (ginsi, unalignlevel 0.6). We manually scrutinized the resulting multiple sequence alignments (File S149, S150) for clear misalignments based on structure-based alignments of available histone domains (File S151, S153). Trees were made using RAxML (version 8.0.20, automatic substitution model selection, GAMMA model of rate heterogeneity, rapid bootstrap analysis of 100 replicates) [12] and/or IQ-TREE (version 1.6.3, extended model selection, ultrafast bootstrap (1000) and SH-like approximate likelihood ratio test) [13]. Trees were visualised and annotated using FigTree [14].

#### Structural similarity and secondary structure prediction

To identify potential homologs based on structural similarity with LECA kinetochore proteins, we searched both the literature and databases such as PFAM (http://pfam.xfam.org [15]), ECOD (http://prodata.swmed.edu/ecod/ [16], RCSB Protein Data Bank (http://rcsb.org [17]) and CATH (http://www.cathdb.info/ [18]). All databases were consulted between January and December 2018. Structures (Table S3) were visualized and processed using the python-based software package Pymol version 2.1.1 [19]. Structural alignments were performed using either 'cealign' and 'super' or were directly downloaded from the aforementioned databases and/or the DALI webserver [20]. The information on the structures and various hyperlinks to databases that we consulted, can be found in Table S3. Note that we did not include our structural analysis in a network-type analysis similar to our profile-versus-profile searches. We chose not to do this as scores of similarity searches do not inform on which are the closest homologs but rather indicate whether a particular protein/domain belongs to a certain structural superfamily. Pymol session files, containing most of the structures used for the comparison of RWD/UBC-like proteins, TBP-like domains and histones are made available (File S151-S153). Secondary structure predictions for Zwint-1 were performed using the JPRED webserver [21], embedded in the alignment package Jalview [22].

#### Classifications and interpretations of homologous protein families

We identified closest homologs of kinetochore proteins based on phylogenetic trees, profile-versus-profile searches and structural similarity. We classified these closest homologs as either eukaryotic or prokaryotic. A closest eukaryotic homolog is a paralog resulting from a gene duplication before LECA [23]. In this, we distinguished kinetochore paralogs from paralogs involved in other eukaryotic cellular processes. If a protein has a kinetochore protein as its closest paralog, likely their ancestral, pre-duplication protein was already part of the primordial kinetochore. Likewise, if more than two kinetochore proteins are most closely related to one another, this group of proteins likely results from multiple successive duplications of a single pre-duplication kinetochore protein. We refer to this ancestral kinetochore protein as ‘anc_KT’ for ‘ancestral kinetochore unit’ (Table 1). Hence, the ‘anc_KT’ is the protein that got involved in the kinetochore, and then duplicated to give rise to the paralogous kinetochore proteins. This ancestral kinetochore protein might have had also closest homolog (either eukaryotic or prokaryotic) outside of the kinetochore, which we also identified. If a LECA kinetochore protein has no closest paralog in the kinetochore, the protein itself forms the ancestral kinetochore unit.

### Supplementary Text

#### Detecting kinetochore homologs using different resources

To complete our picture of the origin of kinetochore proteins, we made use of four different sources of information: phylogenetic trees, (HMM) profile-versus-profile searches and structural information. These different information types result in different qualifications of relationships between pairs of (suspected) homologs (Table 1). In general, the profile-versus-profile searches were in agreement with the relationships observed from the phylogenies (Table S2). However, we noted that various kinetochore proteins, such as RWD proteins and Cep57, hit coiled-coil proteins. In general, the hits we identified as likely coiled-coil were ignored, because coiled-coil similarity might not be indicative of homology, since it could also evolve convergently [24].

#### Proteins in LECA kinetochore & alternative rooting

To determine which proteins were present in the LECA kinetochore (Figure 1B), we first inferred for each protein if it was likely encoded in the genome of LECA. In principle, we did so based on Dollo parsimony, which states that a protein can only be invented once, hence the origin of the protein dates back to the last common ancestor of all species that have it. In applying this approach, we assume that the divergence between Opimoda and Diphoda represents the root of the eukaryotes (Figure S4) [25]. While we are well aware of the controversies about the position of the eukaryotic root [26], we think that alternative rootings would not alter our model of the LECA kinetochore. If for example the root actually lies between (a subset of) Excavata, such as proposed by [27], the presences of kinetochore proteins in this lineage would also support their presence in LECA.

This is the case for the CCAN subunits ('Cenp' proteins, Nkp1 and Nkp2, Figure 1B), because one of the Excavata species (Trichomonas vaginalis) contains CenpX and CenpS [28]. These proteins likely result from duplications, and their closest paralogs are CenpW and CenpT, respectively. Hence, the common ancestor of the Excavata and the other eukaryotes (LECA) likely had these four CCAN components. Furthermore, the recently sequenced genome of the amitochondriate *Monocercomonoides sp. PA203*, an excavate species distantly related to e.g. *Trichomonas vaginalis*, contains four additional CCAN subunits: CenpI, CenpK, CenpL and CenpN (unpublished data). Given that the CCAN subunits strongly co-evolve, likely LECA had the complete CCAN, also under this alternative root. The Dam1 complex would have been inferred to have been present in LECA based on Dollo parsimony, but we think it is very likely that its genes were invented later in evolution and got horizontally transferred among distantly related eukaryotic species [29]. Nkp1 and Nkp2 would not have been found to have been present in LECA under Dollo parsimony. However, we argue that, because they are homologous to subunits of the Mis12 complex (Figure 4A), and because these Mis12 complex subunits are present across the eukaryotic tree of life, Nkp1 and Nkp2 likely resulted from ancient duplications before LECA, giving rise to Nkp1 and Mis12 (Mis12 complex), and to Nkp2 and Nnf1 (Mis12 complex. We infer that Nkp1 and Nkp2 were lost in major eukaryotic lineages quickly after LECA, since we do not observe them in most Diphoda lineages (SAR, Archaeplastida and Excavata, Figure S4). Moreover, Nkp1 and Nkp2 are part of the CCAN, which strongly co-evolves, including Nkp1 and Nkp2 [28].

If a protein likely was encoded by LECA, we in principle hypothesize it to be part of the LECA kinetochore. We nevertheless exclude such a protein from the LECA kinetochore if it depends on a non-LECA protein for the kinetochore function (Hrr25), or if it seems more likely to be involved in another process, as indicated by characterizations in multiple species (Skp1). The complete list of kinetochore proteins studied and considerations for in/excluding them as part of the LECA kinetochore can be found in Table S1.

#### Domain annotation of Mis12-like proteins

In this study, we present the subunits of the Mis12 complex (Mis12, Nnf1, Dsn1, Nsl1) and of the Nkp complex (Nkp1, Nkp2) to be all homologous to one another, having a domain we coin "Mis12-like" (Figure 1, 4A). We inferred their homology using different sources. First, and most convincingly, Nnf1 and Nkp2 appear homologous by being each other's bidirectional best hit in the profile-versus-profile output (Table S2, HHsearch Evalue 10). The Nkp1 and Mis12 full-length profiles hit each each other best with PRC (Table S2, PRC Evalue 10). In the same search, the Nkp2 profile hit the Nnf1 profile, and the Nsl1 profile hit the Mis12 profile. Moreover, the profile of Nkp1 hits that of Mis12 in HHpred online [30], albeit at very high E-value: 180. The structures of the Mis12 subunits were already shown to be similar [31, 32], therefore we propose their homology. If the Mis12 subunits are homologous to one another, and two Mis12 complex subunits are homologous to the two Nkp subunits, all of these six proteins are homologous to each other. Moreover, since we have no indications for other (prokaryotic or eukaryotic) homologs, we infer that the Mis12-like domain was invented before the last eukaryotic common ancestor and gave rise to these six kinetochore proteins via gene duplications.

#### Double RWD domain in Zwint-1 orthologs

In our sensitive profile-versus-profile analysis, various kinetochore RWD proteins hit each other, as well as other RWD-like and E2/UBC proteins, indicating that their sequences were sufficiently similar to confirm their homology, with the notable exceptions of the RWD domains of CenpP (full-length profiles are hit, see Table S2). Interestingly, Zwint-1, the only KMN (Knl1-Mis12-Ndc80) network subunit for which a structure has not yet been determined [33], was hit by various RWD HMM profiles. Indeed, upon further inspection of the predicted secondary structure of Zwint-1 orthologs, we found that they follow a classic tandem RWD topology (Figure S2), similar to its direct interaction partner Knl1 and to the CenpO-CenpP dimer. Interestingly, many Zwint-1 orthologs show degeneration of the C-terminus, resulting in the loss of either the second or both RWD domains; making this a protein family that is particularly hard to predict orthologs for (see also van Hooff et al. [7]). Since all kinetochore RWD proteins form dimers through RWD-RWD interactions and since the main interactor of Zwint-1 is the double RWD protein Knl1, we predict that Zwint-1 is a bona fide double RWD kinetochore protein.

#### RWD evolution

Since bona fide catalytic UBCs were found in both Bacteria and Archaea, ubiquitin-like modification was likely the ancestral function of this fold in FECA [34–37]. Both catalytic [38] and non-catalytic [34] families of the UBC superfamily expanded extensively between FECA and LECA. Functionally, the non-catalytic UBC-like proteins comprise three major groups (see Figure 2): (1) ubiquitin-related vesicle trafficking proteins like Uev1/Tsg101[39] and Aktip [40], (2) a group of E2/E3-related canonical RWD proteins (RWD) involved in DNA/RNA-related processes (e.g. Gcn2 [41] and FancL [42]), and (3) eight LECA kinetochore proteins that form hetero- or homodimers, with either a single RWD: Spc24-Spc25, Mad1-Mad1, Csm1-Csm1, or a double RWD configuration: CenpO-CenpP and Knl1-Zwint-1 (Zwint-1 is also a RWD protein, see ‘Double RWD domain in Zwint-1 orthologs’). Due to the highly divergent sequence evolution of bacterial UBC-like proteins, kinetochore RWD proteins and other non-catalytic UBCs (RWD and for instance Uev1/Tsg101), we could only construct a short alignment (90 positions, minimal 30% column occupancy) for the whole UBC family. Of note, we exclude two E2-like enzymes (Atg3/Atg10) that are involved in autophagy, because they were too divergent to include in the multiple sequence alignment. Both Atg3 and Atg10 are the result of an ancient duplication before LECA [34]. The exact position of these E2s relative to prokaryotic and other eukaryotic UBC-like proteins is unclear. All in all, we performed our analysis using a limited amount of phylogenetic informative positions. Therefore, the parameters for the maximum likelihood methods are potentially overfitted and the resulting phylogenetic tree is therefore likely subject to artefacts such long-branch attraction (see for instance ‘divergent UBC’ and ‘bacteria_E’), resulting in the misplacement of some divergent branches and distortion of the overall tree topology. Nonetheless, overall the eukaryotic UBC superfamily evolved into two distinct groups in eukaryotes: E2 ubiquitin conjugases (UBC, bootstrap:77/100) and two non-catalytic UBC-like groups (RWD and kinetochore RWD; bootstrap:96/100). The inconsistent placement of the second RWD domain of Knl1 and Zwint-1 (Knl1_2 and Zwint-1_2), precluded the conclusive inference for a single origin of kinetochore RWD domains. Whether this means that Knl1_2 and Zwint-1_2 were independently acquired compared to CenpO_2 and CenpP_2 or signify a shared and more complex origin of RWD domains is unclear. Likely the origin of kinetochore RWD domains is closely related to FancL, the only other double RWD protein that is currently found in eukaryotic genomes, and Med15, a mediator complex subunit for which we uncovered the presence of an RWD domain in this study. How bacterial UBCs are related to archaeal and eukaryotic UBCs and RWDs is not entirely clear from our studies. Many different classes of modification systems that operate UBC-like folds are also present in Bacteria (bact_UBC_A-D) and even a non-catalytic domain is found in some lineages (bact_UBC_E) [34]. In addition, a number of bacterial UBC cluster together with archaeal and eukaryotic UBCs. Likely, bacterial UBCs represent an ancient protein modification system (bacteria_UBC_A-E) that has been optimized in archaeal lineage that are closely associated with FECA. Concomitantly, our trees suggest multiple horizontal gene transfers of archaeal/eukaryotic UBCs to bacteria. Various non-catalytic UBC proteins can be found in the Asgard Archaea (closest to the archaeal ancestor of eukaryotes, Figure 1A) [35], which are phylogenetically affiliated with both RWD and UBC (Figure S1E). Therefore, an RWD-like protein could have already been present in the archaeal ancestor of eukaryotes. Given that UBCs of archaeal descent extensively radiated in eukaryotes [38], we think it is likely that RWD and kinetochore RWD were also part of this radiation and are thus of archaeal descent.

#### Reconstruction of histone fold evolution

Although the canonical nucleosomal histones are amongst the most highly conserved eukaryotic proteins at the amino acid level, most other histones in eukaryotes are highly divergent, including the TFIID-related TBP-associated factors (TAF), SAGA-related proteins (SUPT), CCAAT-binding complex/nuclear transcription factor (CBF/NF), Negative coregulator 2 (NC2), subunits of the DNA polymerase epsilon (DPOE), Chromatin Accessibility Complex (CHRAC) and the kinetochore histones (CenpA, CenpS, CenpT, CenpX, CenpW) (see Figure S1I). To produce an informative alignment, we first made separate alignments of slowly evolving orthologs (manually curated) of each of the LECA histone proteins, which we subsequently aligned (corrected based on structural alignments, see ‘Data and Methods’). In addition, we added archaeal and bacterial histone-like sequences, which we acquired through jackhmmer runs against archaeal and bacterial UniProt databases online (see ‘Data and Methods’), using known archaeal histones such as HMf [43], the reported Asgard histone-like proteins [35] and the ‘DUF1931’ protein family as queries. Due to the limited amount of positions (69) and highly divergent nature of the histone family, we could not fully resolve histone evolution as different algorithms (RAxML and IQTREE) and various models gave inconsistent results: (1) a number of different histone groups (H2B, Taf3/8/Supt7, Taf12) appeared at different positions in the tree, (2) the duplication order within for instance the TAF clade was often different with various low bootstrap support values, (3) the position of bacterial and archaeal taxa varied, and (4) the exact placement of CenpX and CenpW relative to each other changed (further detailed below). We here present one of the trees in which CenpX and CenpW are each other’s closest paralog

(Figure S1I). In archaea and bacteria, histones are found that are affliated to different eukaryotic histone groups. The presence of a high number of bacterial histone-like proteins surprised us. Although our analyses did not give a consistent result, it is likely that histone-like proteins in Bacteria were acquired through horizontal gene transfers from either archaeal or eukaryotic lineages. In general, we observe that CenpX and CenpW cluster together with one of two major histone groups: (1) Taf11, H2A, NFY, NC2, DPOE and Chrac1, while CenpS, CenpT and CenpA are more similar to (2) H2B, H3, H4 and all the other TAFs and Supts (not Taf11). The duplication of CenpA and H3 is not always supported and in a number of cases CenpA branches from within the H3 clade. The duplication of CenpT and CenpS is overall well supported (boostrap:87-99/100). The position of CenpX and CenpW varied. The various trees suggested a closest paralog for CenpX, i.e. Taf11, H2A, NC2A and a bacterial clade, and for CenpW, i.e. Taf12 or NC2A/NFY. Apart from the duplication of the kinetochore histone dimer partners, the NFYA-NFYB, NC2A-NC2B and Dpoe3-Doe4/Chrac1 histones seem to originate from an internal duplication as well. The order of duplication for the TFIID and SAGA complex TAF/SUPTs could however not be easily reconciled with the known dimer pairs (see Figure S1I: heterodimers are Taf3/8-Taf10, Taf4-Taf12, Taf6-Taf9, Taf11-Taf13, Supt7-Supt3, Supt3-Taf10 and Ada1-Taf12).

### Supplementary Figures

#### Figure S1. Phylogenetic trees of domains present in kinetochore proteins.

Kinetochore proteins are indicated in green, closest homologs (e.g. eukaryotic closest paralogs) are indicated in blue. Details on tree inference methods can be found in SI, Text, Data and Methods. The sequences were obtained from our local proteome database in combination with bacterial and archaeal entries from the Uniprot database, harvest through online jackhmmer tool (see Data and Methods). The first four letters of the protein name indicate the species as listed in Table S4. Due the size of the tree, the IDs in E (RWD) and I (Histones) are not displayed.

- **(A)** Vps51-like tethering complex subunits (Zw10)
- **(B)** WD40 (Cdc20)
- **(C)** WD40 (Bub3)
- **(D)** kinases (MadBub, Mps1, Aurora, Plk)
- **(E)** RWD (Zwint-1, Knl1, CenpO, CenpP, Spc24, Spc25, Mad1, Csm1)
- **(F)** HORMA (Mad2, p31$^{comet}$)
- **(G)** AAA+ ATPase (Trip13)
- **(H)** NRH-Sec39 (Rod)
- **(I)** histones (CenpA, CenpT, CenpS, CenpX, CenpW) – support is shown for internal nodes that correspond to the origin of new orthologous groups up to the LECA level.

#### Figure S2. Recurrent loss of RWD domains during the evolution of Zwint-1.

**(A)** Secondary structure prediction of three Zwint-1 orthologs: Zwint-1 (human), Kre28 (budding yeast) and Sos7 (fission yeast), reveal a highly divergent C-terminal region. **(B)** Overview of a multiple sequence alignment of 83 Zwint-1 orthologs that was based on the de novo predicted Zwint-like ortholog in zebra fish (*Danio rerio*). The colors of the alignment are based on the classic Clustal coloring scheme and the alignment is condensed so that the letters of the residues are not visible anymore. Note that the second RWD domain is more degenerated and therefore less conserved throughout the eukaryotic lineages that have a Zwint-1 ortholog. On the left the small colored blocks indicate the supergroup to which each species belongs. The red and blue clades indicate which of the orthologs lost the RWD-2 or both the RWD-1 and RWD-2 domain respectively.

**Figure S3. UBC family evolution.**

**(A)** Annotated phylogenetic tree of the UBC superfamily (based on Figure S1E). '{}' denote other domains present and general function for RWD proteins (kinetochore [KT] & other), while for UBC-like proteins it signifies potential substrates. See for SI Text, section *'RWD evolution'* for discussion. **(B)** Evolutionary history of the UBC family: the RWD and UBC proteins in eukaryotes are likely descendant from a bona fide ubiquitin-like modification system [37] that can be found in lineages that are phylogenetically affiliated with the archaeal ancestor of eukaryotes. Subsequent duplications and sub/neofunctionalization gave rise to three groups: (1) UBC (E2 ubiquitin conjugases), (2) RWD ('other', E2/E3-associated proteins), and (3) RWD kinetochore proteins (KT), which are likely the closest homolog to the double RWD protein FancL. The latter two groups are likely highly related. Note the presence of a RWD-like domain in an Asgard protein with 3x RWD/UBC domains. The numbers in light/dark green correspond to the RWD configuration present in each UBC/RWD group (single versus double).

**Figure S4. Phylogeny of eukaryotes.**

Cartoon of the eukaryotic species tree with Opimoda-Diphoda root [2] and eukaryotic supergroups. This topology was used to infer whether a protein was likely present in LECA (Table S1, Figure 1B).

### Supplementary Tables

**Table S1. Kinetochore proteins from human and/or yeast and their inferred presence in the LECA kinetochore (Figure 1B).**

| Protein | Model species | Kinetochore complex | Known domains | Supergroup presence | LECA (y/n: reason) | LECA KT (y/n: reason) |
|---|---|---|---|---|---|---|
| Mad1 | h, y | Mad1-Mad2 | RWD [44] | O, Am, E, S, Ar | yes: parsimony | yes |
| Mad2 | h, y | Mad1-Mad2, MCC | HORMA [45] | O, Am, E, S, Ar | yes: parsimony | yes |
| Bub3 | h, y | MCC | WD40 [46] | O, Am, E, S, Ar | yes: parsimony | yes |
| Cdc20 | h, y | MCC | WD40 [47] | O, Am, E, S, Ar | yes: parsimony | yes |
| MadBub | h, y | MCC | TPR, kinase [48] | O, Am, E, S, Ar | yes: parsimony | yes |
| Mps1 | h, y | | TPR, kinase [49–51] | O, Am, E, S, Ar | yes: parsimony | yes |
| p31 | h | Mad2-Mad2, MCC | HORMA [52] | O, Am, E, S, Ar | yes: parsimony | yes |
| Trip13 | h | Mad2-Mad2, MCC | AAA+ ATPase [53–55] | O, Am, E, S, Ar | yes: parsimony | yes |
| Knl1 | h, y | Knl1-Zwint-1 | RWD (2x) [33] | O, Am, E, S, Ar | yes: parsimony | yes |
| Zwint-1 | h, y | Knl1-Zwint-1 | RWD (2x) [28, 33] | O, Am, E, S, Ar | yes: parsimony | yes |
| Dsn1 | h, y | Mis12-C | Mis12-like [31, 56] | O, Am, E, S, Ar | yes: parsimony | yes |
| Nsl1 | h, y | Mis12-C | Mis12-like [31, 56] | O, Am, S, Ar | yes: parsimony | yes |

| | | | | | | |
|---|---|---|---|---|---|---|
| Nnf1 | h, y | Mis12-C | Mis12-like [31, 32, 56] | O, Am, E, S, Ar | yes: parsimony | yes |
| Mis12 | h, y | Mis12-C | Mis12-like [31, 32, 56] | O, Am, E, S, Ar | yes: parsimony | yes |
| CEP57 | h | | | O, E, S | yes: parsimony | yes |
| ARHGEF17 | h | | RhoGEF [57] | O | no: parsimony | no |
| Ndc80 | h, y | Ndc80-C | CH [58, 59] | O, Am, E, S, Ar | yes: parsimony | yes |
| Spc24 | h, y | Ndc80-C | RWD [60] | O, Am, E, S, Ar | yes: parsimony | yes |
| Spc25 | h, y | Ndc80-C | RWD [60] | O, Am, E, S, Ar | yes: parsimony | yes |
| Nuf2 | h, y | Ndc80-C | CH [58, 59] | O, Am, E, S, Ar | yes: parsimony | yes |
| Ska1 | h | Ska-C | Ska-like [29, 61] | O, Am, S, Ar | yes: parsimony | yes |
| Ska2 | h | Ska-C | Ska-like [[29, 61] | O, Am, E, S, Ar | yes: parsimony | yes |
| Ska3 | h | Ska-C | Ska-like [29, 61] | O, Am, S, Ar | yes: parsimony | yes |
| Ska1 | H | Ska-C | winged-helix [62] | O, Am, S, Ar | yes: parsimony | yes |
| Dam1 | y | Dam1-C | | O, S, Ar | no: HGT [29] | no |
| Duo1 | y | Dam1-C | Duo2-Dad2-like [29] | O, S, Ar | no: HGT [29] | no |
| Dad2 | y | Dam1-C | Duo2-Dad2-like [29] | O, S, Ar | no: HGT [29] | no |
| Dad1 | y | Dam1-C | Dad1/4-Ask1-like [29] | O, Am, S, Ar | no: HGT [29] | no |

| | | | | | | |
|---|---|---|---|---|---|---|
| Dad3 | y | Dam1-C | | O, S, Ar | no: HGT [29] | no |
| Dad4 | y | Dam1-C | Dad1/4-Ask1-like [29] | O, S, Ar | no: HGT [29] | no |
| Hsk3 | y | Dam1-C | | O, S | no: HGT [29] | no |
| Ask1 | y | Dam1-C | Dad1/4-Ask1-like [29] | O, S, Ar | no: HGT [29] | no |
| Spc19 | y | Dam1-C | | O | no: parsimony | no |
| Spc34 | y | Dam1-C | | O, S, Ar | no: HGT [29] | no |
| SKAP | h | SKAP-Astrin | | O | no: parsimony | no |
| Astrin | h | SKAP-Astrin | | O | no: parsimony | no |
| Spindly | h | RZZS | | O | no: parsimony | no |
| Rod | h | RZZS | NRH, Sec39, WD40 [63] | O, E, S | yes: parsimony | yes, but without Spindly, it is unclear how it was recruited to the KT |
| Zwilch | h | RZZS | | O, S | yes: parsimony | yes, but without Spindly, it is unclear how it was recruited to the KT |
| Zw10 | h | RZZS | Vsp51 [64] | O, Am, E, S, Ar | yes: parsimony | yes, but without Spindly, it is unclear how it was recruited to the KT |
| Aurora | h, y | CPC | Kinase [65] | O, Am, E, S, Ar | yes: parsimony | yes |
| Incenp | h, y | CPC | | O, Am, E, S, Ar | yes: parsimony | yes |

| | | | | | | |
|---|---|---|---|---|---|---|
| Survivin | h, y | CPC | Baculoviral IAP repeat (BIR) [66] | O, E, S | yes: parsimony | yes |
| Borealin | h | CPC | | O, Am, E, S, Ar | yes: parsimony | yes |
| Sgo | h, y | CPC | | O, Am, S, Ar | yes: parsimony | yes |
| BugZ | h | | Zinc finger | O, Am, E, S, Ar | yes: parsimony | yes |
| Plk | h, y | | Kinase, polo box [67, 68] | O, Am, E, S, Ar | yes: parsimony | yes |
| CenpA | h, y | Histone | Histone [69] | O, Am, E, S, Ar | yes: parsimony and phylogeny [7] | yes |
| CenpB | h | CCAN | TC5-DDE (TE) [70] | O | no: parsimony | no |
| CenpC | h, y | CCAN | Cupin [71] | O, Am, E, S, Ar | yes: parsimony | yes |
| CenpF | h | | | O | no: parsimony | no |
| CenpE | h | | Kinesin [72] | O, Am, E, S, Ar | yes: parsimony | yes |
| CenpH | h, y | CCAN-HIKM | | O, Am, S | yes: parsimony | yes |
| CenpI | h, y | CCAN-HIKM | HEAT-like repeats [73] | O, Am, S | yes: parsimony | yes |
| CenpK | h, y | CCAN-HIKM | | O, Am, S | yes: parsimony | yes |
| CenpM | h | CCAN-HIKM | GTPase [73] | O, Am, S | yes: parsimony + pre-LECA duplicate | yes |
| CenpL | h, y | CCAN-LN | TBP-like [74] | O, Am, S | yes: parsimony | yes |
| CenpN | h, y | CCAN-LN | TBP-like [74] | O, Am, S | yes: parsimony | yes |

| | | | | | | |
|---|---|---|---|---|---|---|
| CenpO | h, y | CCAN-OPQRU | RWD (2x) [75] | O, Am, S, Ar | yes: parsimony + pre-LECA duplicate | yes |
| CenpP | h, y | CCAN-OPQRU | RWD (2x) [75] | O, Am, S | yes: parsimony + pre-LECA duplicate | yes |
| CenpQ | h, y | CCAN-OPQRU | | O, Am, S | yes: parsimony | yes |
| CenpR | h | CCAN-OPQRU | | O | no: parsimony | no |
| CenpU | h, y | CCAN-OPQRU | | O, Am, S | yes: parsimony | yes |
| Nkp1 | y | CCAN-Nkp | Mis12-like | O | yes: predicted pre-LECA duplicate | yes |
| Nkp2 | y | CCAN-Nkp | Mis12-like | O, Am | yes: predicted pre-LECA duplicate | yes |
| CenpT | h, y | CCAN-TSWX | Histone [76, 77] | O, Am, S | yes: parsimony + pre-LECA duplicate | yes |
| CenpW | h, y | CCAN-TSWX | Histone [76, 77] | O, Am, S | yes: parsimony + pre-LECA duplicate | yes |
| CenpS | h, y | CCAN-TSWX | Histone [77] | O, Am, E, S, Ar | yes: parsimony + pre-LECA duplicate | yes |
| CenpX | h, y | CCAN-TSWX | Histone [77] | O, Am, E, S, Ar | yes: parsimony + pre-LECA duplicate | yes |
| Ndc10 | y | CBF3 | GCR1_C, Crypton F-like [78] | O | no: parsimony | no |
| Ctf13 | y | CBF3 | F-box [79] | O | no: parsimony | no |
| Cep3 | y | CBF3 | Zinc finger, HEAT repeat [80] | O | no: parsimony | no |

| | | | | | | |
|---|---|---|---|---|---|---|
| Skp1 | y | CBF3 | [81] | O, Am, E, S, Ar | yes: parsimony | No: likely operated in SCF ubiquitin ligase complex and was recruited to the KT with the CBF3 complex |
| Csm1 | y | Monopolin | RWD [82] | O, Am, S, Ar | Yes: parsimony | yes |
| Lsr4 | y | Monopolin | [82, 83] | O | No: parsimony | no |
| Mam1 | y | Monopolin | [83] | O | No: parsimony | no |
| Hrr25 | y | Monopolin | Kinase [83] | O, Am, E, S, Ar | Yes: parsimony | No: likely KT function since the origin of Lsr4, Mam1 (Saccharomycetales) |

For each of these proteins, we determined the orthologs across eukaryotic species [7] and determined whether it was encoded by the LECA genome (a ‘LECA protein’), based on Dollo parsimony (present in both Opimoda and Diphoda, Figure S4), or based on the inference of a pre-LECA duplication that gave rise to this protein. In addition, we assessed how likely this protein was part of the LECA kinetochore (‘LECA KT protein’). Model species (h: human, y: budding yeast). Supergroup presence (Opisthokonta (O), Amoebozoa (Am), Excavata (E), SAR (S), Archaeplastida (Ar)).

#### Table S2

Full HHsearch/PRC output (Excel sheets).

#### Table S3

Information on structures analyzed for this study (Excel sheets).

#### Table S4. Species used for phylogenetic analysis (Figure S1)

| Tax ID | Abbreviation | Scientific name | Common name | Eukaryotic supergroup | Group |
|---|---|---|---|---|---|
| 595528 | COWC | Capsaspora owczarzaki ATCC 30864 | Amoeboid symbiont | Opisthokonta | Filasterea |

| | | | | | |
|---|---|---|---|---|---|
| 431895 | MBRE | Monosiga brevicollis MX1 / ATCC 50154 | Choanoflagellate | Opisthokonta | Choanomonada |
| 946362 | SROS | Salpingoeca rosetta | Choanoflagellate | Opisthokonta | Choanomonada |
| 400682 | AQUE | Amphimedon queenslandica | Sponge | Opisthokonta | Metazoa |
| 10228 | TADH | Trichoplax adhaerens | Placozoan | Opisthokonta | Metazoa |
| 45351 | NVEC | Nematostella vectensis | Starlet sea anemone | Opisthokonta | Metazoa |
| 27923 | MLEI | Mnemiopsis leidyi | Sea walnut/Warty comb Jelly | Opisthokonta | Metazoa |
| 6183 | SMAN | Schistosoma mansoni | Flatworm | Opisthokonta | Metazoa |
| 6239 | CELE | Caenorhabditis elegans | Roundworm | Opisthokonta | Metazoa |
| 6279 | BMAL | Brugia malayi | Filarial nematode worm | Opisthokonta | Metazoa |
| 7227 | DMEL | Drosophila melanogaster | Fruitfly | Opisthokonta | Metazoa |
| 180454 | AGAM | Anopheles gambiae str. PEST | Mosquito | Opisthokonta | Metazoa |
| 10224 | SKOW | Saccoglossus kowalevskii | Acorn worm | Opisthokonta | Metazoa |
| 7719 | CINT | Ciona intestinalis | Transparent sea squirt | Opisthokonta | Metazoa |
| 7739 | BFLO | Branchiostoma floridae | Florida lancelet | Opisthokonta | Metazoa |
| 7741 | BBEL | Branchiostoma belcheri | Belcher's lancelet | Opisthokonta | Metazoa |
| 7955 | DRER | Danio rerio | Zebrafish | Opisthokonta | Metazoa |
| 31033 | TRUB | Takifugu rubripes | Japanese pufferfish | Opisthokonta | Metazoa |
| 8364 | XTRO | Xenopus tropicalis | Western clawed frog | Opisthokonta | Metazoa |
| 10090 | MMUS | Mus musculus | Mouse | Opisthokonta | Metazoa |
| 9606 | HSAP | Homo sapiens | Human | Opisthokonta | Metazoa |
| 1993908 | NUSP | Parvularia atlantis | | Opisthokonta | Holomycota |
| 948595 | VCUL | Vavraia culicis floridensis | Microsporidian parasite | Opisthokonta | Microsporidia |
| 876142 | EINT | Encephalitozoon intestinalis ATCC 50506 | Microsporidian parasite | Opisthokonta | Microsporidia |

| | | | | | |
|---|---|---|---|---|---|
| 1003232 | EAED | Edhazardia aedis USNM 41457 | Microsporidian parasite | Opisthokonta | Microsporidia |
| 684364 | BDEN | Batrachochytrium dendrobatidis JAM81 | Chytrid fungus | Opisthokonta | Chytridiomycota |
| 645134 | SPUN | Spizellomyces punctatus DAOM BR117 | Chytrid fungus | Opisthokonta | Chytridiomycota |
| 578462 | AMAC | Allomyces macrogynus ATCC 38327 | Chytrid fungus | Opisthokonta | Blastocladiales |
| 765915 | CANG | Catenaria anguillulae PL171 | Chytrid fungus | Opisthokonta | Blastocladiales |
| 1220926 | MCIR | Mucor circinelloides 1006PhL | Zygomycete fungus | Opisthokonta | Mucoromycotina |
| 763407 | PBLA | Phycomyces blakesleeanus NRRL1555 | Zygomycete fungus | Opisthokonta | Mucoromycotina |
| 1069443 | MVER | Mortierella verticillata NRRL 6337 | Zygomycete fungus | Opisthokonta | Mortierellaceae |
| 310910 | MELO | Mortierella elongata | Zygomycete fungus | Opisthokonta | Mortierellaceae |
| 1357683 | BBER | Basidiobolus meristosporus B9252 | | Opisthokonta | Entomophtorales |
| 796925 | CCOR | Conidiobolus coronatus NRRL28638 | Zygomycete fungus | Opisthokonta | Entomophtorales |
| 763665 | CREV | Coemansia reversa NRRL 1564 | Zygomycete fungus | Opisthokonta | Kickxellomycotina |
| 237631 | UMAY | Ustilago maydis | Fungus | Opisthokonta | Dikarya |
| 235443 | CNEO | Cryptococcus neoformans | Fungus | Opisthokonta | Dikarya |
| 4896 | SPOM | Schizosaccharomyces pombe (strain 972 / ATCC 24843) | Fission yeast | Opisthokonta | Dikarya |
| 367110 | NCRA | Neurospora crassa OR74A | Filamentous fungus | Opisthokonta | Dikarya |
| 284591 | YLIP | Yarrowia lipolytica CLIB 122 | Yeast | Opisthokonta | Dikarya |
| 284592 | DHAN | Debaryomyces hansenii CBS767 | Yeast | Opisthokonta | Dikarya |
| 284590 | KLAC | Kluyveromyces lactis NRRL Y-1140 | Yeast | Opisthokonta | Dikarya |
| 284593 | CGLA | Candida glabrata CBS138 | Yeast | Opisthokonta | Dikarya |
| 559292 | SCER | Saccharomyces cerevisiae S288C | Baker's yeast | Opisthokonta | Dikarya |
| 461836 | TTRA | Thecamonas trahens ATCC 50062 | Bacteriovorous flagellated protist | (sister to Opisthokonta) | Apusomonadida |
| 1257118 | ACAS | Acanthamoebe castellanii str. Neff | Amoeba | Amoebozoa | Discosea |
| 294381 | EHIS | Entamoeba histolytica HM-1:IMSS | Amoeba | Amoebozoa | Entamoebidae |

| | | | | | |
|---|---|---|---|---|---|
| 352472 | DDIS | Dictyostelium discoideum AX4 | Slime mould | Amoebozoa | Dictyosteliida |
| 670386 | PPAL | Polysphondylium pallidum PN500 | Cellular slime mold | Amoebozoa | Dictyosteliida |
| 1410327 | ASUB | Acytostelium subglobosum LB1 | | Amoebozoa | Dictyosteliida |
| 280463 | EHUX | Emiliania huxleyi CCMP1516 | Haptophyte | (sister to SAR) | Haptophyta |
| 1460289 | CTOB | Chrysochromulina tobin strain CCMP291 | Haptophyte | (sister to SAR) | Haptophyta |
| 453998 | MONO | Monocercomonoides sp. PA203 | | Excavata | Oxymonadida |
| 412133 | TVAG | Trichomonas vaginalis G3 | Protozoan | Excavata | Parabasalia |
| 941442 | GINT | Giardia intestinalis assemblage A | Protozoan | Excavata | Fornicata |
| 347515 | LMAJ | Leishmania major strain Friedlin | Protozoan | Excavata | Discicristata |
| 185431 | TBRU | Trypanosoma brucei TREU 927 | Protozoan | Excavata | Discicristata |
| 3039 | EGRA | Euglena gracilis | | Excavata | Discicristata |
| 744533 | NGRU | Naegleria gruberi strain NEG-M | Protozoan | Excavata | Discicristata |
| 12968 | BHOM | Blastocystis hominis | Single-celled protozoan parasite | SAR | Stramenopiles |
| 702273 | AKER | Aplanochytrium kerguelense PBS07 | Marine protist | SAR | Stramenopiles |
| 717989 | ALIM | Aurantiochytrium limacinum ATCC MYA-1381 | Marine protist | SAR | Stramenopiles |
| | SPAR | | | | |
| 403677 | PINF | Phytophthora infestans T30-4 | Potato late blight fungus | SAR | Stramenopiles |
| 890382 | ALAI | Albugo laibachii Nc14 | Arabidopsis pathogen | SAR | Stramenopiles |
| 272952 | HPAR | Hyaloperonospora parasitica | former name: Peronospora parasitica. Also called Hyaloperonospora arabidopsidis | SAR | Stramenopiles |
| 1093141 | NGAD | Nannochloropsis gaditana CCMP526 | Stramenopile alga | SAR | Stramenopiles |
| 44056 | AANO | Aureococcus anophagefferens CCMP1984 | Harmful bloom alga | SAR | Stramenopiles |
| 2880 | ESIL | Ectocarpus siliculosus | Brown alga | SAR | Stramenopiles |

| | | | | | |
|---|---|---|---|---|---|
| 309737 | COKA | Cladosiphon okamuranus | Brown alga | SAR | Stramenopiles |
| 556484 | PTRI | Phaeodactylum tricornutum CCAP1055/1 | Diatome | SAR | Stremenopiles |
| 296543 | TPSE | Thalassiosira pseudonana | Marine diatom | SAR | Stremenopiles |
| 423536 | PMAR | Perkinsus marinus ATCC 50983 | | SAR | Alveolata |
| 1202447 | SMIN | Symbiodinium minutum | Dinoflagellate | SAR | Alveolata |
| 36329 | PFAL | Plasmodium falciparum 3D7 | Protozoan | SAR | Alveolata |
| 414452 | CPAI | Cryptosporidium parvum Iowa II | Protozoan | SAR | Alveolata |
| 508771 | TGON | Toxoplasma gondii ME49 | Protozoan | SAR | Alveolata |
| 5888 | PTET | Paramecium tetraurelia | Paramecium | SAR | Alveolata |
| 5911 | TTHE | Tetrahymena thermophila | Tetrahymena | SAR | Alveolata |
| 1172189 | OTRI | Oxytricha trifallax | Ciliate | SAR | Alveolata |
| 5963 | SCOE | Stentor coeruleus | Ciliate | SAR | Alveolata |
| 753081 | BNAT | Bigelowiella natans CCMP2755 | Chlorarachniophyte alga | SAR | Rhizaria |
| 905079 | GTHE | Guillardia theta CCMP2712 | Cryptomonad alga | Archaeplastida | Cryptophyceae |
| 2762 | CPAR | Cyanophora paradoxa | Glaucophyte | Archaeplastida | Glaucophyta |
| 130081 | GSUL | Galdieria sulphuraria | Red alga | Archaeplastida | Rhodophyceae |
| 280699 | CMER | Cyanidioschyzon merolae 10D | Red alga | Archaeplastida | Rhodophyceae |
| 554065 | CVAR | Chlorella variabilis NC64A | Green alga | Archaeplastida | Chloroplastida |
| 574566 | CSUB | Coccomyxa subellipsoidea C-169 | Green alga | Archaeplastida | Chloroplastida |
| 3055 | CREI | Chlamydomonas reinhardtii | Green alga | Archaeplastida | Chloroplastida |
| 3068 | VCAR | Volvox carteri | Green alga | Archaeplastida | Chloroplastida |
| 296587 | MSPE | Micromonas species RCC299 | Picoplanktonic green alga | Archaeplastida | Chloroplastida |
| 436017 | OLUC | Ostreococcus lucimarinus CCE9901 | Green alga | Archaeplastida | Chloroplastida |
| 1075084 | BPRA | Bathycoccus prasinos RCC1005 | Green alga | Archaeplastida | Chloroplastida |

| | | | | | |
|---|---|---|---|---|---|
| 3175 | KFLA | Klebsormidium flaccidum | | Archaeplastida | Klebsormidiaceae |
| 145481 | PPAT | Physcomitrella patens subsp. patens | Moss | Archaeplastida | Chloroplastida |
| 88036 | SMOE | Selaginella moellendorffii | Spikemoss | Archaeplastida | Chloroplastida |
| 13333 | ATRI | Amborella trichopoda | | Archaeplastida | Chloroplastida |
| 39947 | OSAT | Oryza sativa japonica | Rice | Archaeplastida | Chloroplastida |
| 3702 | ATHA | Arabidopsis thaliana | Thale cress | Archaeplastida | Chloroplastida |
| 218851 | ACOE | Aquilegia coerulea Goldsmith | Rocky mountain columbine | Archaeplastida | Chloroplastida |

#### Supplementary Files

HMM profiles used in this study (1-147)

Cytoscape files of the HHsearch network (148)

Alignments of RWD/UBC-like (149) and histones (150) used for phylogenetic analysis

Pymol session files of RWD-like/UBC (151), TBP-like (152), histones (153)

Prokaryotic operon HORMA-Trip13 overview linked to Figure 4B (154)

### Supplementary References

1. Katoh K, Standley DM (2013) MAFFT multiple sequence alignment software version 7: improvements in performance and usability. Mol Biol Evol 30:772–780 . doi: 10.1093/molbev/mst01
2. Mi H, Huang X, Muruganujan A, et al (2017) PANTHER version 11: expanded annotation data from Gene Ontology and Reactome pathways, and data analysis tool enhancements. Nucleic Acids Res 45:183–189 . doi: 10.1093/nar/gkw1138
3. Madera M (2008) Profile Comparer: a program for scoring and aligning profile hidden Markov models. Bioinformatics 24:2630–2631 . doi: 10.1093/bioinformatics/btn504
4. Söding J (2005) Protein homology detection by HMM-HMM comparison. Bioinformatics 21:951–960 . doi: 10.1093/bioinformatics/bti125
5. Shannon P, Markiel A, Ozier O, et al (2003) Cytoscape: a software environment for integrated models of biomolecular interaction networks. Genome Res 13:2498–2504
6. Velankar S, Dana JM, Jacobsen J, et al (2013) SIFTS: Structure Integration with Function, Taxonomy and Sequences resource. Nucleic Acids Res 41:483–489 . doi: 10.1093/nar/gks1258
7. van Hooff JJ, Tromer E, van Wijk LM, et al (2017) Evolutionary dynamics of the kinetochore network in eukaryotes as revealed by comparative genomics. EMBO Rep 18:1559–1571 . doi: 10.15252/embr.201744102
8. van Wijk Berend LM; S (2018) Phylogenomics reveals ancestral kinase relations, repertoire, and fate in present-day eukaryotes. Prep
9. Burroughs AM, Zhang D, Schäffer DE, et al (2015) Comparative genomic analyses reveal a vast, novel network of nucleotide-centric systems in biological conflicts, immunity and signaling. Nucleic Acids Res 43:10633–10654 . doi: 10.1093/nar/gkv1267
10. Katoh K (2002) MAFFT: a novel method for rapid multiple sequence alignment based on fast Fourier transform. Nucleic Acids Res 30:3059–3066 . doi: 10.1093/nar/gkf436
11. Capella-Gutiérrez S, Silla-Martínez JM, Gabaldón T (2009) trimAl: a tool for automated alignment trimming in large-scale phylogenetic analyses. Bioinformatics 25:1972–1973 . doi: 10.1093/bioinformatics/btp348
12. Stamatakis A (2014) RAxML version 8: a tool for phylogenetic analysis and post-analysis of large phylogenies. Bioinformatics 30:1312–1313 . doi: 10.1093/bioinformatics/btu033
13. Nguyen L-T, Schmidt HA, von Haeseler A, Minh BQ (2015) IQ-TREE: A Fast and Effective Stochastic Algorithm for Estimating Maximum-Likelihood Phylogenies. Mol Biol Evol 32:268–274 . doi: 10.1093/molbev/msu300
14. Rambaut A (2012) FigTree v1. 4. Mol Evol phylogenetics Epidemiol Edinburgh, UK Univ Edinburgh, Inst Evol Biol
15. Finn RD, Bateman A, Clements J, et al (2014) Pfam: The protein families database. Nucleic Acids Res. 42:222–230
16. Cheng H, Schaeffer RD, Liao Y, et al (2014) ECOD: An Evolutionary Classification of Protein Domains. PLoS Comput Biol 10:e1003926 . doi: 10.1371/journal.pcbi.1003926
17. Berman HM, Westbrook J, Feng Z, et al (2000) The Protein Data Bank. Nucleic Acids Res 28:235–242 . doi: 10.1093/nar/28.1.235
18. Dawson NL, Lewis TE, Das S, et al (2017) CATH: An expanded resource to predict protein function through structure and sequence. Nucleic Acids Res 45:289–295 . doi: 10.1093/nar/gkw1098
19. Schrödinger L (2010) PyMOL The PyMOL Molecular Graphics System
20. Holm L, Sander C (1995) Dali: a network tool for protein structure comparison. Trends Biochem Sci 20:478–480 . doi: https://doi.org/10.1016/S0968-0004(00)89105-7
21. Drozdetskiy A, Cole C, Procter J, Barton GJ (2015) JPred4: a protein secondary structure prediction server. Nucleic Acids Res 43:389–394 . doi: 10.1093/nar/gkv332

22. Waterhouse AM, Procter JB, Martin DMA, et al (2009) Jalview Version 2—a multiple sequence alignment editor and analysis workbench. Bioinformatics 25:1189–1191 . doi: 10.1093/bioinformatics/btp033
23. Makarova KS, Wolf YI, Mekhedov SL, et al (2005) Ancestral paralogs and pseudoparalogs and their role in the emergence of the eukaryotic cell. Nucleic Acids Res 33:4626–4638 . doi: 10.1093/nar/gki775
24. Mistry J, Finn RD, Eddy SR, et al (2013) Challenges in homology search: HMMER3 and convergent evolution of coiled-coil regions. Nucleic Acids Res 41:e121 . doi: 10.1093/nar/gkt263
25. Derelle R, Torruella G, Klimeš V, et al (2015) Bacterial proteins pinpoint a single eukaryotic root. Proc Natl Acad Sci U S A 112:693–699 . doi: 10.1073/pnas.1420657112
26. Williams TA (2014) Evolution: Rooting the eukaryotic tree of life. Curr. Biol. 24:151–152
27. He D, Fiz-Palacios O, Fu C-J, et al (2014) An Alternative Root for the Eukaryote Tree of Life. Curr Biol 24:465–470 . doi: 10.1016/j.cub.2014.01.036
28. Tromer EC (2017) Evolution of the Kinetochore Network in Eukaryotes. Utr Univ. doi: 10.13140/RG.2.2.16846.56640
29. van Hooff JJE, Snel B, Kops GJPL (2017) Unique Phylogenetic Distributions of the Ska and Dam1 Complexes Support Functional Analogy and Suggest Multiple Parallel Displacements of Ska by Dam1. Genome Biol Evol 9:1295–1303 . doi: 10.1093/gbe/evx088
30. Zimmermann L, Stephens A, Nam S-Z, et al (2018) A Completely Reimplemented MPI Bioinformatics Toolkit with a New HHpred Server at its Core. J Mol Biol 430:2237–2243 . doi: 10.1016/j.jmb.2017.12.007
31. Petrovic A, Keller J, Liu Y, et al (2016) Structure of the MIS12 Complex and Molecular Basis of Its Interaction with CENP-C at Human Kinetochores. Cell 167:1028–1040 . doi: 10.1016/j.cell.2016.10.005
32. Zhou X, Zheng F, Wang C, et al (2017) Phosphorylation of CENP-C by Aurora B facilitates kinetochore attachment error correction in mitosis. Proc Natl Acad Sci 114:10677–10676 . doi: 10.1073/pnas.1710506114
33. Petrovic A, Mosalaganti S, Keller J, et al (2014) Modular Assembly of RWD Domains on the Mis12 Complex Underlies Outer Kinetochore Organization. Mol Cell 53:591–605 . doi: 10.1016/j.molcel.2014.01.019
34. Burroughs AM, Jaffee M, Iyer LM, Aravind L (2008) Anatomy of the E2 ligase fold: implications for enzymology and evolution of ubiquitin/Ub-like protein conjugation. J Struct Biol 162:205–218 . doi: 10.1016/j.jsb.2007.12.006
35. Zaremba-Niedzwiedzka K, Caceres EF, Saw JH, et al (2017) Asgard archaea illuminate the origin of eukaryotic cellular complexity. Nature 541:353–358 . doi: 10.1038/nature21031
36. Nunoura T, Takaki Y, Kakuta J, et al (2011) Insights into the evolution of Archaea and eukaryotic protein modifier systems revealed by the genome of a novel archaeal group. Nucleic Acids Res 39:3204–3223 . doi: 10.1093/nar/gkq1228
37. Hennell James R, Caceres EF, Escasinas A, et al (2017) Functional reconstruction of a eukaryotic-like E1/E2/(RING) E3 ubiquitylation cascade from an uncultured archaeon. Nat Commun 8:1120 . doi: 10.1038/s41467-017-01162-7
38. Grau-Bove X, Sebe-Pedros A, Ruiz-Trillo I (2015) The eukaryotic ancestor had a complex ubiquitin signaling system of archaeal origin. Mol Biol Evol 32:726–739 . doi: 10.1093/molbev/msu334
39. Sundquist WI, Schubert HL, Kelly BN, et al (2004) Ubiquitin Recognition by the Human TSG101 Protein. Mol Cell 13:783–789 . doi: 10.1016/S1097-2765(04)00129-7
40. Xu L, Sowa ME, Chen J, et al (2008) An FTS/Hook/p107(FHIP) complex interacts with and promotes endosomal clustering by the homotypic vacuolar protein sorting complex. Mol Biol Cell 19:5059–5071 . doi: 10.1091/mbc.E08-05-0473
41. Nameki N, Yoneyama M, Koshiba S, et al (2004) Solution structure of the RWD domain of the mouse GCN2 protein. Protein Sci 13:2089–2100 . doi: 10.1110/ps.04751804

42. Hodson C, Cole AR, Lewis LPC, et al (2011) Structural analysis of human FANCL, the E3 ligase in the Fanconi anemia pathway. J Biol Chem 286:32628–32637 . doi: 10.1074/jbc.M111.244632
43. Mattiroli F, Bhattacharyya S, Dyer PN, et al (2017) Structure of histone-based chromatin in Archaea. Science 357:609–612 . doi: 10.1126/science.aaj1849
44. Kim S, Sun H, Tomchick DR, et al (2012) Structure of human Mad1 C-terminal domain reveals its involvement in kinetochore targeting. Proc Natl Acad Sci 109:6549–6554 . doi: 10.1073/pnas.1118210109
45. Aravind L, Koonin E V (1998) The HORMA domain: a common structural denominator in mitotic checkpoints, chromosome synapsis and DNA repair. Trends Biochem Sci 23:284–286 . doi: 10.1016/S0968-0004(98)01257-2
46. Wilson DK, Cerna D, Chew E (2005) The 1.1-Å Structure of the Spindle Checkpoint Protein Bub3p Reveals Functional Regions. J Biol Chem 280:13944–13951 . doi: 10.1074/jbc.M412919200
47. Tian W, Li B, Warrington R, et al (2012) Structural analysis of human Cdc20 supports multisite degron recognition by APC/C. Proc Natl Acad Sci 109:18419
48. Bolanos-Garcia VM, Blundell TL (2011) BUB1 and BUBR1: multifaceted kinases of the cell cycle. Trends Biochem Sci 36:141–150 . doi: 10.1016/j.tibs.2010.08.004
49. Lee S, Thebault P, Freschi L, et al (2012) Characterization of spindle checkpoint kinase mps1 reveals domain with functional and structural similarities to tetratricopeptide repeat motifs of Bub1 and BubR1 checkpoint kinases. J Biol Chem 287:5988–6001 . doi: 10.1074/jbc.M111.307355
50. Thebault P, Chirgadze D, Dou Z, et al (2012) Structural and functional insights into the role of the N-terminal Mps1 TPR domain in the spindle assembly checkpoint (SAC). Biochem J. doi: 10.1042/BJ20121448
51. Nijenhuis W, von Castelmur E, Littler D, et al (2013) A TPR domain–containing N-terminal module of MPS1 is required for its kinetochore localization by Aurora B. J Cell Biol 201:217–231 . doi: 10.1083/jcb.201210033
52. Yang M, Li B, Tomchick DR, et al (2007) p31comet Blocks Mad2 Activation through Structural Mimicry. Cell 131:744–755 . doi: https://doi.org/10.1016/j.cell.2007.08.048
53. Ye Q, Kim DH, Dereli I, et al (2017) The AAA+ ATPase TRIP13 remodels HORMA domains through N-terminal engagement and unfolding. EMBO J 36:2419–2434 . doi: 10.15252/embj.201797291
54. Ye Q, Rosenberg SC, Moeller A, et al (2015) TRIP13 is a protein-remodeling AAA+ ATPase that catalyzes MAD2 conformation switching. Elife 4:e07367
55. Brulotte ML, Jeong B-C, Li F, et al (2017) Mechanistic insight into TRIP13-catalyzed Mad2 structural transition and spindle checkpoint silencing. Nat Commun 8:1956 . doi: 10.1038/s41467-017-02012-2
56. Dimitrova YN, Jenni S, Valverde R, et al (2016) Structure of the MIND Complex Defines a Regulatory Focus for Yeast Kinetochore Assembly. Cell 167:1014–1027 . doi: 10.1016/j.cell.2016.10.011
57. Rümenapp U, Freichel-Blomquist A, Wittinghofer B, et al (2002) A mammalian Rho-specific guanine-nucleotide exchange factor (p164-RhoGEF) without a pleckstrin homology domain. Biochem J 366:721–728 . doi: 10.1042/BJ20020654
58. Ciferri C, Pasqualato S, Screpanti E, et al (2008) Implications for kinetochore-microtubule attachment from the structure of an engineered Ndc80 complex. Cell 133:427–439
59. Schou KB, Andersen JS, Pedersen LB (2014) A divergent calponin homology (NN-CH) domain defines a novel family: Implications for evolution of ciliary IFT complex B proteins. Bioinformatics 30:899–902 . doi: 10.1093/bioinformatics/btt661
60. Wei RR, Schnell JR, Larsen NA, et al (2006) Structure of a central component of the yeast kinetochore: the Spc24p/Spc25p globular domain. Structure 14:1003–1009 . doi: 10.1016/j.str.2006.04.007
61. Jeyaprakash AA, Santamaria A, Jayachandran U, et al (2012) Structural and functional organization of the Ska complex, a key component of the kinetochore-microtubule interface. Mol Cell 46:274–286 . doi: 10.1016/j.molcel.2012.03.005

62. Abad MA, Medina B, Santamaria A, et al (2014) Structural basis for microtubule recognition by the human kinetochore Ska complex. Nat Commun 5:2964 . doi: 10.1038/ncomms3964
63. Çivril F, Wehenkel A, Giorgi FM, et al (2010) Structural analysis of the RZZ complex reveals common ancestry with multisubunit vesicle tethering machinery. Structure 18:616–626 . doi: 10.1016/j.str.2010.02.014
64. Tripathi A, Ren Y, Jeffrey PD, Hughson FM (2009) Structural characterization of Tip20p and Dsl1p, subunits of the Dsl1p vesicle tethering complex. Nat Struct &Amp; Mol Biol 16:114 . doi: 10.1038/nsmb.1548https://www.nature.com/articles/nsmb.1548#supplementary-information
65. Sessa F, Mapelli M, Ciferri C, et al (2005) Mechanism of Aurora B Activation by INCENP and Inhibition by Hesperadin. Curr Biol 18:379–391
66. Jeyaprakash AA, Klein UR, Lindner D, et al (2007) Structure of a Survivin–Borealin–INCENP Core Complex Reveals How Chromosomal Passengers Travel Together. Cell 131:271–285 . doi: https://doi.org/10.1016/j.cell.2007.07.045
67. Elling RA, Fucini R V, Romanowski MJ (2008) Structures of the wild-type and activated catalytic domains of Brachydanio rerio Polo-like kinase 1 (Plk1): changes in the active-site conformation and interactions with ligands. Acta Crystallogr Sect D 64:909–918 . doi: doi:10.1107/S0907444908019513
68. Cheng K, Lowe ED, Sinclair J, et al (2003) The crystal structure of the human polo-like kinase-1 polo box domain and its phospho-peptide complex. EMBO J 22:5757–5768 . doi: 10.1093/emboj/cdg558
69. Tachiwana H, Kagawa W, Shiga T, et al (2011) Crystal structure of the human centromeric nucleosome containing CENP-A. Nature 476:232 . doi: 10.1038/nature10258https://www.nature.com/articles/nature10258#supplementary-information
70. Kipling D, Warburton PE (1997) Centromeres, CENP-B and Tigger too. Trends Genet 13:141–145 . doi: https://doi.org/10.1016/S0168-9525(97)01098-6
71. Cohen RL, Espelin CW, De Wulf P, et al (2008) Structural and functional dissection of Mif2p, a conserved DNA-binding kinetochore protein. Mol Biol Cell 19:4480–4491 . doi: 10.1091/mbc.E08-03-0297
72. Garcia-Saez I, Yen T, Wade RH, Kozielski F (2004) Crystal Structure of the Motor Domain of the Human Kinetochore Protein CENP-E. J Mol Biol 340:1107–1116 . doi: https://doi.org/10.1016/j.jmb.2004.05.053
73. Basilico F, Maffini S, Weir JR, et al (2014) The pseudo GTPase CENP-M drives human kinetochore assembly. Elife 3:e02978
74. Pentakota S, Zhou K, Smith C, et al (2017) Decoding the centromeric nucleosome through CENP-N. Elife 6: . doi: 10.7554/eLife.33442
75. Schmitzberger F, Harrison SC (2012) RWD domain: a recurring module in kinetochore architecture shown by a Ctf19–Mcm21 complex structure. EMBO Rep 13:216–222 . doi: 10.1038/embor.2012.1
76. Hori T, Amano M, Suzuki A, et al (2008) CCAN Makes Multiple Contacts with Centromeric DNA to Provide Distinct Pathways to the Outer Kinetochore. Cell 135:1039–1052 . doi: https://doi.org/10.1016/j.cell.2008.10.019
77. Nishino T, Takeuchi K, Gascoigne KE, et al (2012) CENP-T-W-S-X forms a unique centromeric chromatin structure with a histone-like fold. Cell 148:487–501 . doi: 10.1016/j.cell.2011.11.061
78. Cho U-S, Harrison SC (2011) Ndc10 is a platform for inner kinetochore assembly in budding yeast. Nat Struct &Amp; Mol Biol 19:48 . doi: 10.1038/nsmb.2178https://www.nature.com/articles/nsmb.2178#supplementary-information
79. Russell ID, Grancell AS, Sorger PK (1999) The Unstable F-box Protein p58-Ctf13 Forms the Structural Core of the CBF3 Kinetochore Complex. J Cell Biol 145:933–950 . doi: 10.1083/jcb.145.5.933
80. Bellizzi JJ, Sorger PK, Harrison SC (2007) Crystal Structure of the Yeast Inner Kinetochore Subunit Cep3p. Structure 15:1422–1430 . doi: https://doi.org/10.1016/j.str.2007.09.008
81. Leber V, Nans A, Singleton MR (2018) Structural basis for assembly of the CBF3 kinetochore complex.

EMBO J 37:269–281 . doi: 10.15252/embj.201798134
82. Corbett KD, Yip CK, Ee LS, et al (2010) The monopolin complex crosslinks kinetochore components to regulate chromosome-microtubule attachments. Cell 142:556–567 . doi: 10.1016/j.cell.2010.07.017
83. Ye Q, Ur SN, Su TY, Corbett KD (2016) Structure of the Saccharomyces cerevisiae Hrr25:Mam1 monopolin subcomplex reveals a novel kinase regulator. EMBO J 35:2139–2151 . doi: 10.15252/embj.201694082
